# GluD1 Modulates GluN2B-containing NMDAR Function and Plasticity at Subicular Synapses

**DOI:** 10.64898/2026.08.11.744000

**Authors:** Eric Purisic, Darielle Lewis-Sanders, Marian Zhong, Joshua Stamos, Tianhua Wang, Christian Valade, Markus Wöhr, Eric Sobie, Jinye Dai

## Abstract

Dysregulation of the delta-type glutamate receptor GluD1 and N-methyl-D-aspartate receptors (NMDARs) is implicated in neuropsychiatric disorders including schizophrenia and intellectual disability, and GluD1 modulates NMDAR response in hippocampal neurons. However, the precise mechanisms by which GluD1 influences specific NMDAR subtypes remain undefined, representing a critical gap given the reliance of synaptic plasticity and cognition on NMDAR composition. GluN2A- and GluN2B-containing NMDARs are essential for synaptic long-term potentiation (LTP) and contextual learning and memory. Here, we used CRISPR/Cas9 to generate GluD1 knockout (KO) in cultured hippocampal neurons and observed a selective decrease in GluN2B-containing NMDAR responses. In acute hippocampal slices, GluD1 KO similarly reduced GluN2B-containing NMDAR currents at ventral CA1→subiculum synapses and impaired LTP at these synapses. In vivo, region-specific GluD1 deficiency in the ventral subiculum disrupted long-term contextual memory, indicating a critical role for GluD1 in cognitive processes. These findings demonstrate that GluD1 is indispensable for preserving GluN2B-containing NMDAR function, synaptic plasticity, and memory, providing molecular insight into how GluD1 regulates NMDAR subtypes implicated in synaptic dysfunction in neuropsychiatric disorders. Understanding this mechanism will guide the development of therapeutic strategies that selectively target GluD1-dependent modulation of NMDAR subtypes in brain disease.

## INTRODUCTION

Synaptic communication underlies neuronal network functions, and ionotropic glutamate receptors (iGluRs) are central mediators of excitatory transmission and plasticity. iGluRs include AMPA receptors (AMPARs), NMDA receptors (NMDARs), kainate receptors (GluKs), and delta-type receptors (GluDs). Although GluD receptors (GluD1 and GluD2) were identified decades ago, their atypical properties—such as the absence of conventional glutamate binding—led to early underestimation of their physiological roles (*1–7*). Emerging genetic and functional evidence, however, implicates GluD receptors in critical aspects of brain function (*8–15*).

GluD1 (encoded by *GRID1*) is broadly expressed at postsynaptic sites across diverse brain regions. Human genetic studies link *GRID1* variants to neurodevelopmental and neuropsychiatric disorders, including intellectual disability, schizophrenia, and Rett syndrome, each featuring cognitive impairments (*16–22*). Mouse models show that GluD1 global deletion disrupts cognitive functions such as fear memory (*23, 24*). Together, these studies underscore that GluD1 contributes to cognition, yet the underlying synaptic mechanisms and molecular pathways remain poorly defined.

At the cellular level, GluDs localize to dendritic spines and influence excitatory and inhibitory synapse formation and maturation (*8–10, 14, 15, 25–29*). Its homolog GluD2 has well-established roles in cerebellar synaptic plasticity (e.g., long-term depression in Purkinje cells) (*9, 10, 30, 31*), but the physiological roles of GluD1 in forebrain circuits are less clear. Recent work indicates that GluD1 regulates excitatory synapse development in the hippocampus and medial prefrontal cortex and modulates NMDAR function in hippocampal neurons (*28, 32–39*). Importantly, distinct NMDAR subunit compositions critically shape synaptic properties: GluN2B (NMDAR subunit 2B)-containing NMDA receptors (GluN2B-containing NMDARs) predominate early in development, and progressive incorporation of GluN2A (NMDAR subunit 2A)-containing NMDA receptors (GluN2A-containing NMDARs) with maturation alters channel kinetics and plasticity thresholds (*40–44*). In addition, studies have identified three major synaptic NMDAR assemblies in mature cortical and hippocampal neurons: diheteromeric GluN1/GluN2A and GluN1/GluN2B receptors, as well as triheteromeric GluN1/GluN2A/GluN2B receptors, each estimated to comprise approximately one-third of the total NMDAR population (*43, 45*). In hippocampal neurons, both GluN2A and GluN2B contribute to synaptic plasticity, but they do so with distinct temporal profiles and downstream signaling roles (*46–52*). Despite this, it remains unknown which specific NMDAR subtypes are engaged by GluD1 and how subtype-selective regulation by GluD1 influences synaptic maturation and plasticity. Thus, determining whether GluD1 preferentially controls GluN2B-containing or GluN2A-containing NMDARs is essential for understanding its contribution to developmental synaptic remodeling and cognitive processes.

Here, we first examined developmental expression patterns and found that GluD1, GluN2A, and GluN2B mRNA levels increase in parallel from postnatal day 0 (P0) to P21 *in vivo* and from days *in vitro* 1 (DIV1) to DIV9 in cultured hippocampal neurons, then stabilize by adulthood (P60 or DIV16), accompanied by a decrease in the GluN2B/GluN2A ratio (Figures 1 and 3). Using CRISPR/Cas9-mediated GluD1 KO at an early developmental timepoint (DIV4 in cultures and P21-23 in brains) in hippocampal neurons, we observed reduced NMDAR-mediated excitatory postsynaptic current (EPSC) amplitude and faster decay kinetics in both cultured neurons and acute slices, indicating a selective loss of GluN2B-containing NMDAR-mediated responses at the synaptic surface (Figures 1 and 3). Supporting this interpretation, application of the GluN2B-selective antagonist Ro 25-6981 (1 µM) produced a similar reduction in EPSC amplitude and acceleration of decay kinetics in control neurons but had no additional effect in GluD1 KO neurons. Notably, at this low concentration, Ro 25-6981 preferentially inhibits diheteromeric GluN1/GluN2B receptors while minimally affecting triheteromeric GluN1/GluN2A/GluN2B receptors due to their distinct pharmacological properties (*53–58*). Accordingly, the approximately 35-40% reduction in NMDAR-mediated EPSC amplitude following GluD1 deletion is consistent with previous estimates of the contribution of diheteromeric GluN1/GluN2B receptors to the total NMDAR population in mature neurons (Figures 1 and 3) (*45, 52, 59–61*). Furthermore, the selective reduction in synaptic surface GluN1 and GluN2B staining provides additional support for a loss of GluN2B-containing NMDARs in GluD1 KO neurons (Figure 2). These results reveal a previously unrecognized role for GluD1 in maintaining GluN2B-containing NMDARs signaling during synapse maturation.

**Figure 1.**
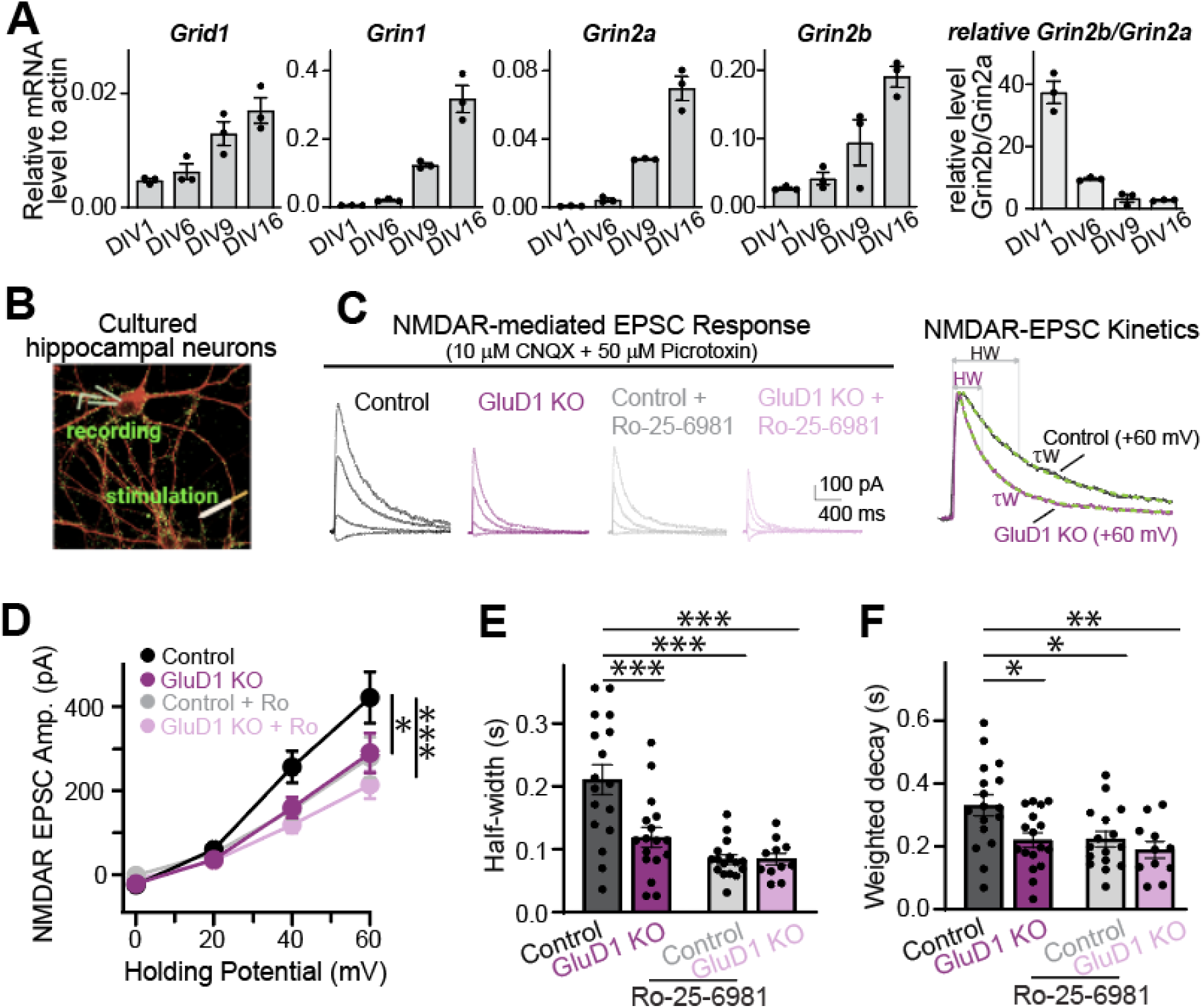
GluD1 Regulates GluN2B-NMDAR-Mediated Synaptic Transmission in Cultured Hippocampal Neurons. (A) Summary graph showing mRNA levels (normalized for β-actin levels) as measured by qRT-PCR of cultured hippocampal neurons at DIV 1, 4, 9, 16 from Cas9 mice. (B) NMDAR-EPSC recordings from CRISPR/Cas9 GluD1 KO and control hippocampal cultures at DIV15-17. Configuration of evoked NMDAR-EPSCs recordings (with 10 μM CNQX and 50 μM picrotoxin), measured at holding potential of 0 mV, +20 mV, +40 mV, and +60 mV. We utilized Cas9 cultured neurons with lentiviral or AAV infections of CRISPR sgRNAs to delete GluD1 as previous study (*34, 105*). (C) Left, the sample traces of current-voltage (I-V) relationship of NMDAR-EPSCs in control, KO, control with Ro-25-6981, and KO with Ro-25-6981 conditions. Right, sample NMDAR-EPSC traces of +60 mV from control and GluD1 KO, displaying width (HW) and weighted decay time constant (τw) calculated using double exponential function fitting: A(t) =A_slow_exp(−t/t_slow_) +A_fast_exp(−t/t_fast_), where t_slow_ and t_fast_ are the decay time constants of the slow and fast component, and A_slow_ and A_fast_ are their respective amplitudes. To compare the decay of EPSCs at different conditions, the weighted time constant (τw) will be used: τw = t_slow_[A_slow_/(A_slow_+ A_fast_)] + t_fast_[A_fast_/(A_slow_+ A_fast_)]. (D-F) Summary graphs of amplitude (D), HW (E), and τw (F) in control (n=17/5), KO (n=18/5), control with Ro-25-6981 (n=16/3), and KO with Ro-25-6981 (n=11/3) conditions. Two-way/one-way ANOVA test with Turkey Post-hoc shows significant differences. Numbers of neurons/mice (*n*) are shown. Data are mean ± s.e.m.

**Figure 2.**
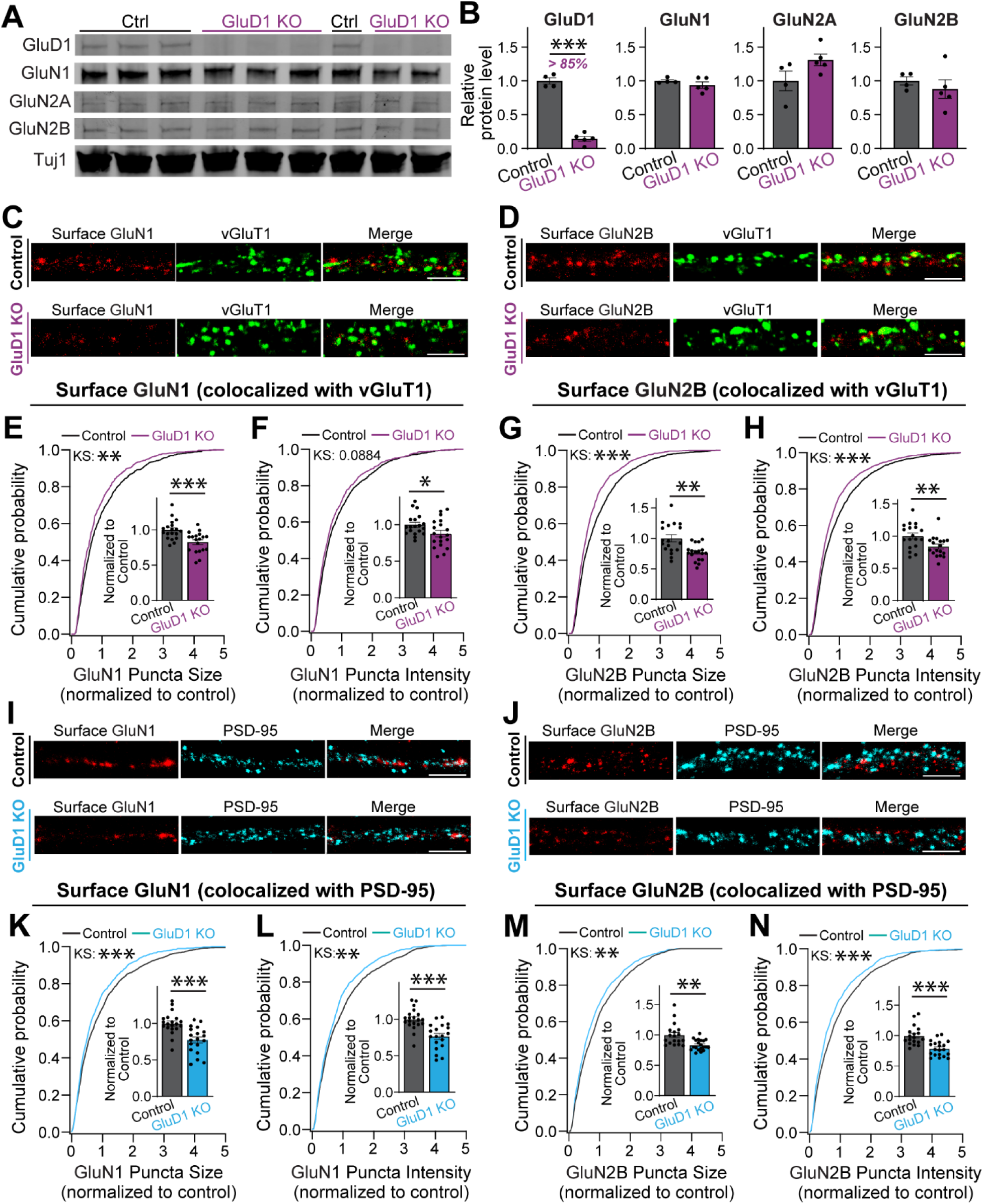
GluD1 KO Does Not Alter Total NMDAR Subunit Expression But Selectively Reduces Synaptic Surface Expression. (A) Representative immunostaining blot showing protein expression levels of GluD1, GluN1, GluN2A, GluN2B, and Tuj1 from DIV17 cultured hippocampal neurons. We utilized Cas9 neurons with AAV infections of CRISPR sgRNA to KO GluD1. (B) Quantification of relative protein expression levels in GluD1 condition (n = 5, normalized to control conditions after normalized to Tuj1) compared to the control condition (n = 4). (C and D) Synaptic surface staining and confocal imaging of vGluT1 and NMDA subunits in cultured hippocampal neurons. Representative images of dendrites in Control and GluD1 KO conditions. bar: 5 μm. (E-H) Bar graphs and cumulative probability graphs of normalized puncta size and intensity for GluN1 (E and F, n = 19 per condition) and GluN2B (G and H, n = 18 per condition) from 3 batches of coverslips per condition. Unpaired two-tailed t-test shows significant difference in normalized GluN1 and GluN2B puncta size and intensity. Kolmogorov-Smirnov test was performed for cumulative probability graphs. (I and J) Synaptic surface staining and confocal imaging of PSD-95 and NMDA subunits in cultured hippocampal neurons. Representative images of dendrites in Control and GluD1 KO conditions. bar: 5 μm. (K-N) Bar graphs and cumulative probability graphs of normalized puncta size and intensity for GluN1 (K, n = 20 and L, n = 19) and GluN2B (M, n = 18 and N, n = 20) from 3 batches of coverslips per condition. Unpaired two-tailed t-test shows significant difference in normalized GluN1 and GluN2B puncta size and intensity. Kolmogorov-Smirnov test was performed for cumulative probability graphs. Unpaired two-tailed t-test or Mann-Whitney U test shows significant differences. Numbers of neurons/mice (*n*) are shown. Data are mean ± s.e.m.

**Figure 3.**
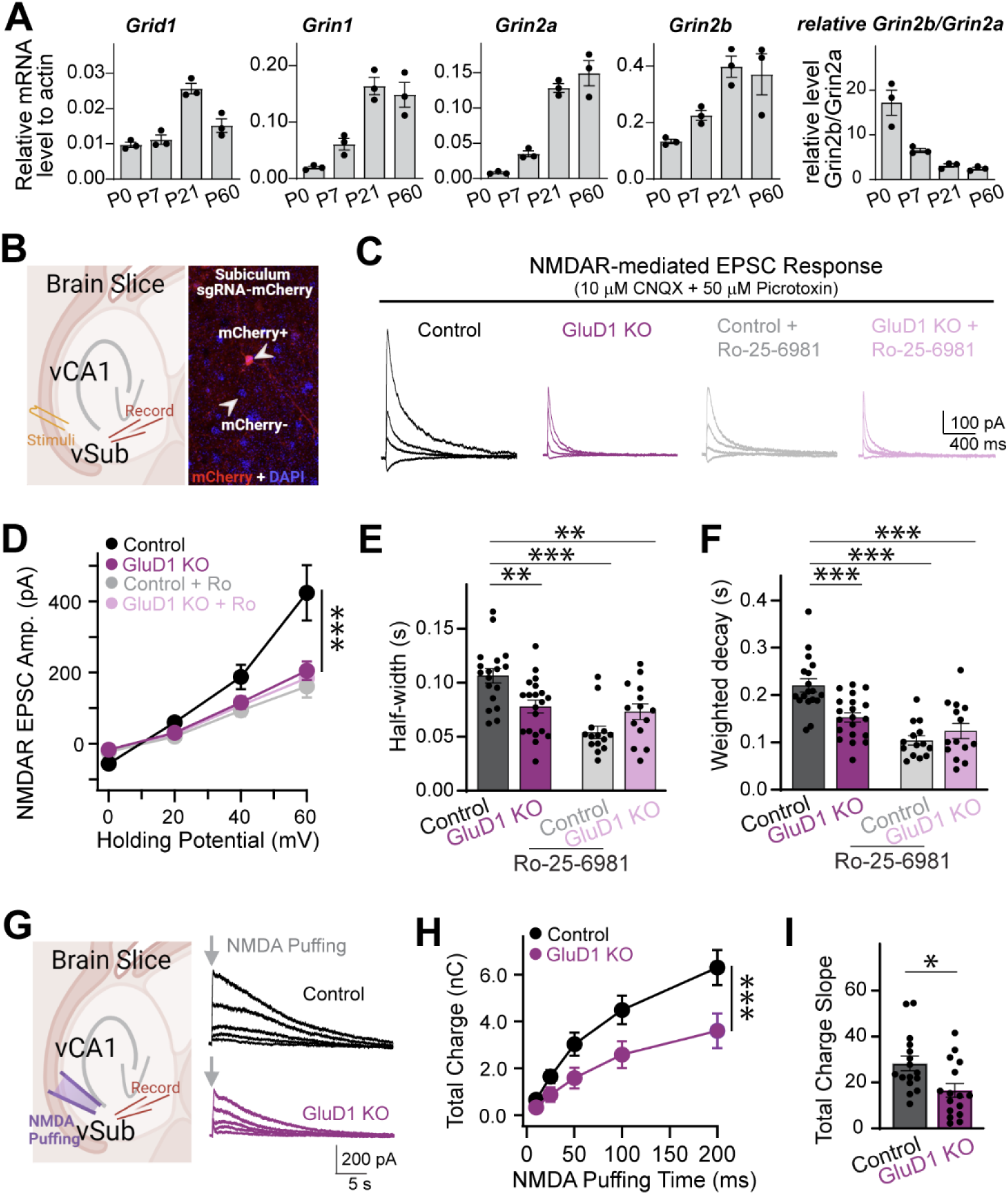
GluD1 Regulates GluN2B-NMDAR-Mediated Synaptic Transmission in Hippocampal vCA1→vSub Synapses. (A) Summary graph showing mRNA levels (normalized for β-actin levels) as measured by qRT-PCR of hippocampal tissues from P0, 7, 21, 60 of Cas9 mice. (B) CRISPR-mediated GluD1 KO and I-V curves of NMDAR-EPSCs in vCA1→vSub synapses of Cas9 KI mice. Left, configuration of evoked NMDAR-EPSCs recordings in vCA1→vSub synapses (with 10 μM CNQX and 50 μM picrotoxin). Right, confocal image of subiculum region indicates mCherry positive (mCherry+) and mCherry negative (mCherry-) neurons. Brain slices were obtained from 6-9 weeks old mice that received lentiviral sgRNA-mCherry or diluted AAV-sgRNA-tdTomato injections at P21-23. (C) Sample EPSC traces were measured at holding potential of 0 mV, +20 mV, +40 mV, and +60 mV in control, GluD1 KO, control with Ro-25-6981, and KO with Ro-25-6981 conditions. (D-F) Summary graphs of amplitude (D), HW (E), and τw (F) in control (n = 18), KO (n = 20), control with Ro-25-6981 (n = 14), and KO with Ro-25-6981 (n = 14) conditions from ≥ 3 mice. (G-I) Postsynaptic NMDAR-mediated currents were recorded from patched subicular pyramidal neurons in response to local NMDA application. NMDA was puffed at 50 μM in the presence of picrotoxin and CNQX using a picospritzer, with the puffing pipette positioned approximately two cell-body distances from the recorded neuron. NMDA was applied with increasing puff durations of 10, 25, 50, 100, and 200 ms. Representative traces are shown in G. Summary graphs show NMDA-evoked total charge transfer (H) and slope (I) in control (tdTomato negative, n = 16) and GluD1 KO neurons (tdTomato positive, n = 16) from 3 mice. Two-way/one-way ANOVA test with Turkey Post-hoc or Kruskal-Wallis test with Dunn’s Post-hoc multiple comparison or unpaired two-tailed Mann-Whitney U test shows significant differences. Data are mean ± s.e.m.

Since GluN2B-containing NMDAR is involved in the GluD1 signaling pathway, we sought to understand the extent to which GluD1 influences synaptic plasticity across hippocampal circuits. Although one study reported no change in LTP at CA3-CA1 synapses after GluD1 knockdown (*28*), the diversity of hippocampal outputs, such as the CA1→subiculum pathway, calls for broader investigation. We focused on ventral CA1→subiculum (vCA1→vSub) synaptic plasticity, given the high GluD1 expression in subiculum, its role as a major hippocampal output, and its relevance for emotion-related memory (*11, 62–66*). Subicular pyramidal neurons comprise regular-firing and burst-firing types, each with distinct long-term potentiation (LTP) mechanisms (postsynaptic NMDAR-dependent LTP involving changes in AMPAR levels in regular-firing neurons; presynaptic LTP involving changes in release probability in burst-firing neurons) (*65, 67–70*). We found that GluD1 deletion selectively impairs NMDAR-dependent LTP in regular-firing vSub neurons, identifying a cell-type-specific requirement for GluD1 in hippocampal plasticity (Figure 4). Moreover, region-specific deletion of GluD1 in the vSub impairs long-term contextual fear memory, indicating that GluD1 in this output region is necessary for proper retrieval of aversive memories (Figure 5). This finding aligns with the known role of the vCA1→vSub circuit in memory processing and highlights the behavioral relevance of GluD1-mediated synaptic plasticity in this region.

**Figure 4.**
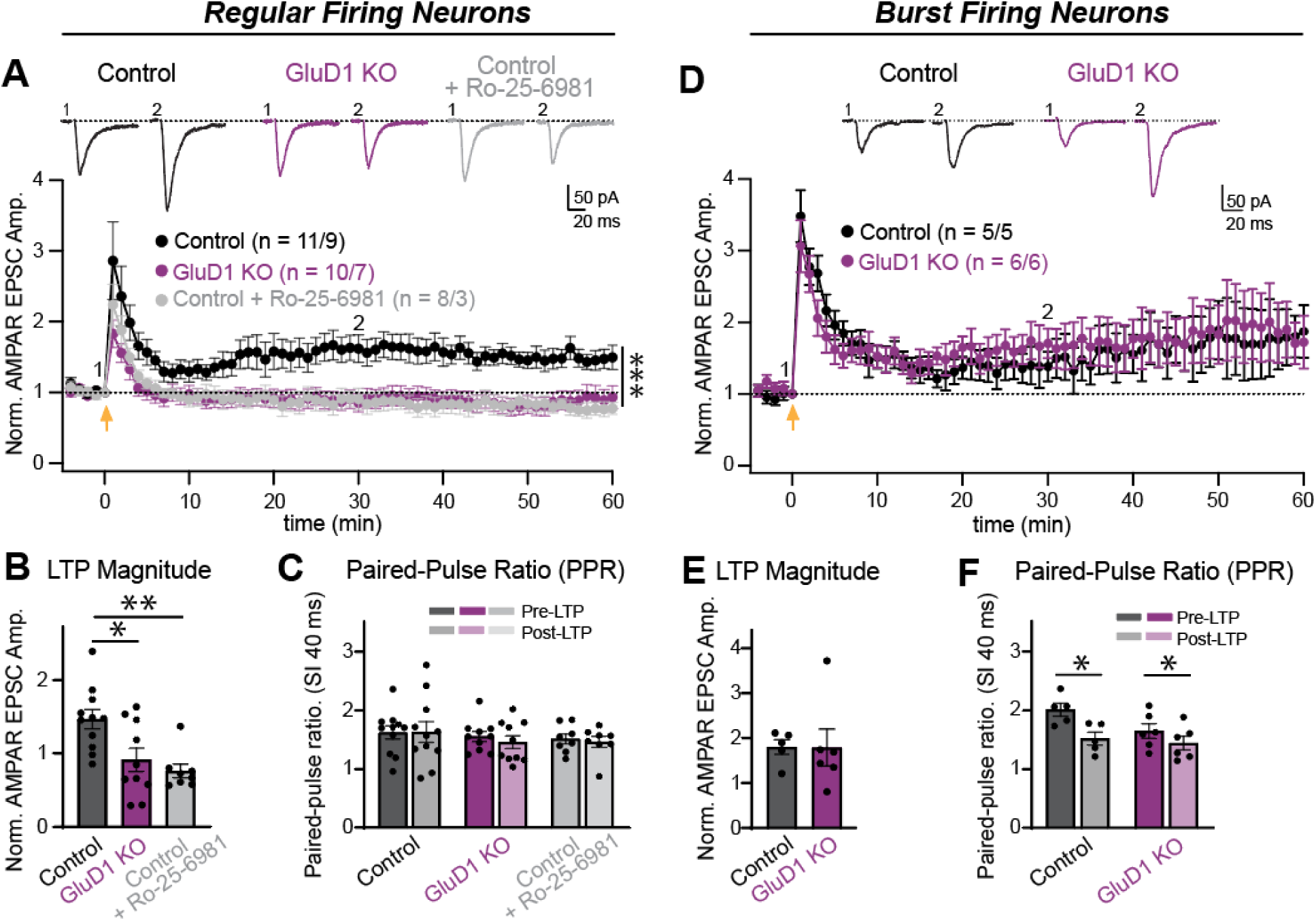
CRISPR-mediated GluD1 KO or Blocking GluN2B-containing NMDARs inhibits LTP in the Regular Subicular Firing Neurons. (A) Recordings from regular-spiking neurons in 6-9-week-old Cas9 knock-in mice showed that a high-frequency protocol (4 × 1-s trains at 100 Hz, 10-s inter-train interval; delivered at the yellow arrow) induced robust LTP under wild-type conditions but not after GluD1 KO or blocking GluN2B-containing NMDARs. (B and C) Summary graphs (B) depict the LTP magnitude calculated as the normalized average EPSC amplitudes during the last 5 minutes after LTP induction. Kruskal-Wallis test with Dunn’s Post-hoc multiple comparison shows significant difference. Summary graphs of PPR (C) with stimulus interval (SI 40 ms) before and after LTP. Paired two-tailed t-test or Mann-Whitney U test shows no significant difference. (D) Recordings from burst-spiking neurons in 6-9-week-old Cas9 knock-in mice showed that a high-frequency protocol (4 × 1-s trains at 100 Hz, 10-s inter-train interval; delivered at the yellow arrow) induced robust LTP under both wild-type and GluD1 conditions. (E and F) Summary graphs (E) depict the LTP magnitude calculated as the normalized average EPSC amplitudes during the last 5 minutes after LTP induction. Unpaired two-tailed Mann-Whitney U test shows no significant difference. Summary graphs of PPR (F) with stimulus interval (SI 40 ms) before and after LTP. Paired two-tailed t-test shows significant difference. Numbers of neurons/mice (*n*) are shown. Data are mean ± s.e.m.0

**Figure 5.**
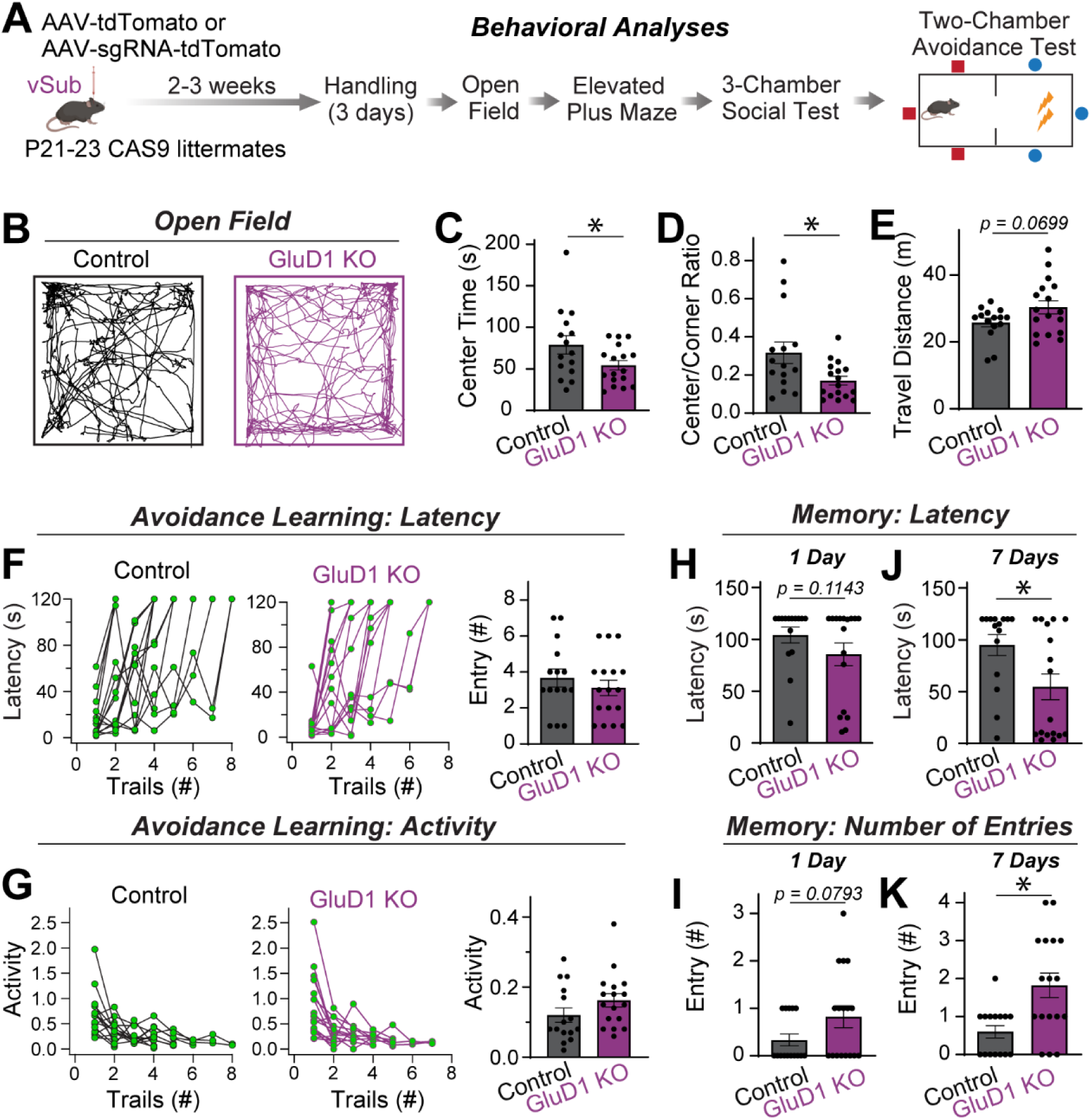
The Role of GluD1 In vSub Controls Anxiety-Like Behavior and Contextual Memory. (A) Strategy of experimental design. The vSub region of Cas9 mice was infected using stereotactic injections at P21-23 with AAVs expressing tdTomato (retains WT genotype) or sgRNA-tdTomato (produces GluD1 KO genotype), and mice were analyzed after 2-3 weeks infection. (B-E) Open field test. Example tracking of control and GluD1 KO in vSub (B). Summary graphs of center time (C), center/corner ratio (D), and total travel distance (E) in 10 mins for control (n=15) and GluD1 KO (n=17) mice. Two-tailed t-test shows significant differences in center time and center/corner ratio. (F and G) Two-chamber avoidance test. Deletion of GluD1 in the hippocampal vSub did not affect avoidance learning. (F) Trial-by-trial latency during training in control and GluD1 KO mice (left and middle) and the number of chamber entries (right). (G) Trial-by-trial locomotor activity in the safe chamber (left and middle) and average activity during the final training trial (right). (H-K) GluD1 controls long-term contextual memory in mice as measured by conditioned avoidance. Entry latency and number of entries of mice on day 1 and day 7 after training were measured. Unpaired one-tailed Mann-Whitney U test shows significance between GluD1 WT (n=15) and GluD1 KO (n=17) mice in day 7 long-term contextual memory. Data are mean ± s.e.m.

Together, these findings establish that GluD1 is critical for preserving GluN2B-containing NMDAR-mediated synaptic signaling during maturation and for enabling NMDAR-dependent LTP in defined hippocampal circuits. Elucidating the molecular interactions by which GluD1 controls the synaptic surface component of GluN2B-containing NMDARs and synaptic potentiation will deepen our understanding of synaptic maturation mechanisms and illuminate how GluD1 dysfunction contributes to cognitive deficits across brain disorders.

## RESULTS

### CRISPR-mediated GluD1 Knockout Accelerates Kinetics of NMDAR Responses in Hippocampal Synapses

In cultured hippocampal neurons, we found parallel increases in Grid1, Grin1, Grin2a, and Grin2b mRNA, encoding GluD1, GluN1, GluN2A, and GluN2B respectively, from days *in vitro* 1 (DIV1) to DIV16 and decreased relative ratio of Grin2b/Grin2a (Figure 1A). In prior work, we showed that GluD1 is essential for NMDAR-mediated synaptic transmission (*34*), but the NMDAR subtypes involved remained undefined. To address this, we used a CRISPR/Cas9 approach with either Lenti or AAV expressing single guide RNA (sgRNA) at DIV 4 to delete GluD1 in cultured hippocampal neurons, confirming efficient KO at the protein level at DIV 15-17 (Figure 2A and Figure S1). For functional readout at DIV 15-17, whole-cell voltage-clamp recordings were then performed at depolarized holding potentials (+0, +20, +40, +60 mV) to enhance NMDAR activation while minimizing the Mg²⁺ blockade. Synaptic NMDAR-EPSCs were isolated by bath application of CNQX to block AMPAR/GluK currents and picrotoxin to block GABAR currents (Figure 1B and 1C). In GluD1 KO neurons, evoked NMDAR-EPSC amplitude was significantly reduced compared to control neurons, indicating an overall loss of functional synaptic NMDARs (Figure 1D). Unexpectedly, analysis of decay kinetics revealed a substantial acceleration: both the half-width (the time between EPSC onset and decay to 50% of peak) and the weighted decay time constant τw (calculated from double-exponential fits as (A₁τ₁ + A₂ τ₂)/(A₁ + A₂)) were markedly decreased in GluD1 KO neurons (Figure 1E and 1F). Because GluN2A-containing and GluN2B-containing NMDARs exhibit distinct decay profiles, with GluN2A-mediated currents decaying rapidly, whereas GluN2B-mediated currents decaying more slowly, the faster kinetics in GluD1 KO neurons suggested a selective reduction of the slower GluN2B-containing NMDAR component.

To test whether the kinetic changes reflected the loss of GluN2B-containing NMDARs, we applied the GluN2B-selective antagonist Ro 25-6981 (1 µM). At this low concentration, Ro 25-6981 preferentially blocks diheteromeric GluN1/GluN2B receptors while minimally affecting triheteromeric GluN1/GluN2A/GluN2B receptors (*53–57*). In control neurons, GluN2B blockade recapitulated the GluD1 KO phenotype: NMDAR-EPSC amplitude decreased by a similar fraction, and both half-width and τw were shortened to the same extent observed in GluD1-deficient cells (Figure 1B-1F). In contrast, applying the same antagonist in GluD1 KO neurons produced no further reduction in amplitude or change in kinetics, indicating occlusion (Figure 1B-1F). In addition, the ∼40% reduction in NMDAR-mediated EPSC amplitude observed after either GluD1 deletion or Ro-25-6981 blockade closely corresponds to the estimated proportion of diheteromeric GluN1/GluN2B receptors to the total NMDAR population in mature neurons (Figure 1D) (*45, 52, 60, 61*). Together with the accelerated NMDAR-EPSC decay kinetics after GluD1 deletion or Ro-25-6981 blockade, these findings are suggestive of a preferential reduction in diheteromeric GluN1/GluN2B receptor function. However, we cannot fully exclude the possibility that triheteromeric GluN1/GluN2A/GluN2B receptors are also affected. Thus, our results support a model in which GluD1 maintains functional GluN2B-containing NMDAR signaling at hippocampal excitatory synapses, with the available pharmacological and kinetic evidence pointing toward an effect on diheteromeric GluN1/GluN2B receptors.

### GluD1 KO Reduces Surface Expression of GluN1 and GluN2B at Excitatory Hippocampal Synapses

To determine whether the functional deficits in NMDAR-mediated synaptic transmission after GluD1 KO reflect altered total NMDAR expression or perturbed surface trafficking, we first quantified overall protein levels of GluN1, GluN2A, and GluN2B by Western blot using whole-cell lysis of cultured hippocampal neurons at DIV 15-17 (Figure 2A). Despite the clear reduction in NMDAR-EPSC amplitude and altered kinetics in GluD1 KO neurons, total levels of GluN1, GluN2A, and GluN2B were unchanged relative to controls (Figure 2B), suggesting that GluD1 does not affect global expression of these subunits. We therefore hypothesized that GluD1 specifically influences the synaptic surface stabilization of GluN2B-containing NMDARs.

To test this hypothesis, we performed immunocytochemistry to label extracellular epitopes of the NMDAR subunits: GluN1, GluN2A, and GluN2B. Cultured neurons were first stained with subunit-specific antibodies under non-permeabilizing conditions, then permeabilized and co-stained for presynaptic vesicular glutamate transporter 1 (vGluT1) to label excitatory synapses. At vGluT1-positive presynaptic sites, quantitative puncta analysis showed a significant reduction in both the average size and integrated intensity of surface GluN1 puncta and surface GluN2B puncta in GluD1 KO neurons (Figure 2C-2H). In contrast, vGluT1 puncta and surface GluN2A puncta were unchanged (Figure S2). Cumulative probability analyses further confirmed these selective shifts in surface GluN1 and GluN2B puncta, but not in surface GluN2A puncta (Figure 2C-2H and Figure S2C-S2D).

To independently assess synaptic surface NMDAR subunits using a postsynaptic marker, we next quantified surface GluN1 and GluN2B puncta colocalized with postsynaptic density protein 95 (PSD-95). GluD1 KO significantly reduced the average size and integrated intensity of PSD-95-colocalized surface GluN1 and GluN2B puncta (Figure 2I-2N). In contrast, PSD-95 puncta size and intensity were unchanged in GluD1 KO (Figure S2B). Cumulative probability distributions further confirmed the reductions in surface GluN1 and GluN2B puncta (Figure 2K-2N).

These results strongly support a model in which GluD1 regulates the synaptic surface expression of GluN2B-containing NMDARs, without altering overall subunit abundance. The reduction of surface GluN1 puncta indicates the decrease in total NMDAR-mediated synaptic currents, while the selective loss of GluN2B at synapses accounts for the accelerated NMDAR-EPSC kinetics in GluD1 KO neurons. Collectively, these data indicate that GluD1 is required to maintain GluN2B-containing NMDARs at the synaptic surface site, thereby modulating both the magnitude and kinetic profile of NMDAR-mediated synaptic transmission.

### GluD1 KO Reduces NMDAR-EPSCs and Accelerates Receptor Kinetics at vCA1**→**vSub Synapses

Our findings in cultured hippocampal neurons establish a clear link between GluD1 and GluN2B-containing NMDARs. However, it remains unknown whether GluD1 similarly modulates GluN2B-NMDAR subtypes under physiological conditions *in vivo*. We found that GluD1 mRNA expression reaches a peak at P21 and then stabilizes through adulthood at P60 (Figure 3A). Thus, we decided to delete GluD1 at P21-23 in vSub to determine if GluD1 also plays a crucial role in contributing to the maintenance of GluN2B-NMDAR responses in vCA1→vSub hippocampal synapses during synaptic maturation *in vivo*.

To this aim, we used AAV infection of the vSub at P21-23 and conducted *ex vivo* brain slice electrophysiology at the age of 6-9 weeks (Figure 3B). Our findings indicate a consistent reduction phenotype following GluD1 KO in vCA1→vSub synapses (Figure 3C and 3D), which aligns with observations made in cultured hippocampal synapses (Figure 1). Furthermore, the kinetics of NMDAR-EPSCs accelerate in GluD1 KO subicular neurons, indicating a reduction in the GluN2B-containing NMDAR subtypes (Figure 3E and 3F). To confirm this reduction is due to GluN2B-containing NMDARs, we once again employed pharmacological blockade of GluN2B-containing NMDARs using 1 μM Ro-25-6981 to mimic potential functional changes seen in GluD1 KO neurons. Our results show that blockade of GluN2B-containing NMDARs in control mimics the phenotypes observed in GluD1 KO, including a reduction in NMDAR-mediated EPSC amplitude and a reduction in half-width and weighted decay (Figure 3B-3F). Blockade in the GluD1 KO does not show a significant difference, suggesting that GluD1 KO is sufficient to elicit the reduction of GluN2B-containing NMDARs (Figure 3B-3F). Overall, our findings illustrate that GluD1 KO a) reduces overall NMDAR-mediated synaptic transmission at vCA1→vSub synapses; b) accelerates the NMDAR kinetics; c) mimics pharmacological blockade of GluN2B-containing NMDARs. This result is consistent with our findings in cultured hippocampal neurons. Specifically, the ∼35% reduction in NMDAR-mediated EPSC amplitude following GluD1 deletion is comparable to the estimated contribution of diheteromeric GluN1/GluN2B receptors to the total NMDAR population in mature neurons (Figure 3) (*45, 52, 59–61*). Together with the accelerated decay kinetics and the pharmacological results, these data support the conclusion that GluD1 substantially modulates GluN2B-containing NMDAR-mediated synaptic transmission at vCA1→vSub synapses, potentially through a preferential effect on diheteromeric GluN1/GluN2B NMDAR signaling.

To determine whether the reduced synaptic NMDAR response reflected a broader loss of NMDARs from the neuronal surface, we next measured whole-cell responses to exogenous NMDA application in acute brain slices. NMDA was locally puffed onto neurons for five stimulus durations, allowing us to assess surface NMDAR function across a range of activation periods. GluD1 deletion produced a significant reduction in the total charge transfer of NMDA-evoked currents (Figure 3G and 3H). Consistently, the relationship between charge transfer and puff duration showed a reduced slope in GluD1-deficient neurons (Figure 3I). Together, these findings support a role for GluD1 in maintaining functional NMDARs at the neuronal surface.

### GluD1 KO Impairs LTP Induction at vCA1**→**vSub Synapses

Since GluN2B-containing NMDARs are highly involved in synaptic plasticity (*46–52*), we next examined the connection between GluD1 and GluN2B-containing NMDARs in regulating LTP at vCA1→vSub synapses of 6-9 week-old mice after deletion of GluD1 at P21-23. These synapses on regular- and burst-firing subiculum neurons exhibit distinct forms of LTP, with the former expressing a postsynaptic NMDAR-dependent form of LTP, whereas the latter displays a presynaptic form of LTP (*38, 69, 70*). Employing the same timeline of GluD1 deletion, we found that the GluD1 deletion did not affect presynaptic LTP in burst-firing neuron synapses, which undergo a characteristic change in paired-pulse ratios (PPRs) after LTP induction (Figure 4D-4F). However, the GluD1 deletion abolished postsynaptic LTP in regular-firing neurons without a change of PPRs after induction (Figure 4A-4C). Thus, the GluD1 deletion selectively ablates NMDAR-dependent postsynaptic LTP in regular-firing subiculum neurons without affecting presynaptic LTP in burst-firing neurons. Our previous experiments showed that GluD1 KO primarily reduces GluN2B-containing NMDARs at the synaptic surface; therefore, inhibition of LTP in regular-firing subiculum neurons due to GluD1 deletion may be through a GluN2B-containing NMDAR-mediated mechanism. To test this hypothesis, we employed the GluN2B-selective antagonist, Ro-25-6981, in the control condition.

Interestingly, GluN2B-blocking in control reproduces the phenotype observed with GluD1 KO alone, showing the same changes in LTP magnitude (Figure 4A-4C). This suggests that the impaired LTP in regular-firing subicular GluD1 KO neurons is due to the reduction of GluN2B-containing NMDARs at these synapses. Together, this novel finding represents a potential mechanism for GluD1 in modulating synaptic plasticity at vCA1→vSub synapses through an GluN2B-containing NMDAR-mediated mechanism.

### GluD1 KO in Ventral Subiculum Impairs Long-Term Contextual Memory in a Two-Chamber Avoidance Task

So far, our data showed that GluD1 KO clearly impairs synaptic plasticity at vCA1→vSub synapses through its effects on GluN2B-containing NMDARs. However, the behavioral significance of these observed changes is unclear. To follow up on these findings and to examine behavioral phenotypes associated with GluD1 KO in the vSub, we performed a battery of behavioral tests (Figure 5A). Given the diversity of behavioral phenotypes associated with both vSub and GluD1 function, we included a broad set of tests to capture potential anxiety- and depression-like behaviors as well as cognitive and social deficits. Our findings show a prominent reduction in center time and a trend toward increased distance traveled in the open field test (Figure 5B-5E). This indicates a potential anxiety-like and hyperactive phenotype in vSub GluD1 KO mice. However, we observed no differences in the elevated plus maze or three-chamber sociability test as compared to control mice (Figure S3 and S4).

To relate our observations on synaptic plasticity at vCA1→vSub synapses to cognitive functioning, we next assessed effects on learning and memory using a well-established two-chamber avoidance task (*38, 39*). We observed no clear deficits during the learning phase, as indicated by the lack of difference in entries into the foot-shock chamber during training, suggesting intact memory acquisition (Figure 5F and 5G). However, one week after training, GluD1 KO mice showed a reduced latency to enter and an increased number of entries into the foot-shock chamber, as compared to control mice. In fact, the first evidence for impaired memory was already obtained one day after training, with GluD1 KO mice displaying a tendency toward reduced latency and altered entry numbers (Figure 5H-5K). These findings indicate that GluD1 KO does not impair learning, such as fear memory acquisition associated with this two-chamber avoidance task, but instead leads to deficits in long-term contextual fear memory retrieval.

## DISCUSSION

Our study demonstrates that the orphan receptor GluD1 is a critical regulator of GluN2B-containing NMDAR responses and synaptic plasticity, with effects on long-term contextual fear memory. By combining measurements in cultured hippocampal neurons and vCA1→vSub synapses, we show that loss of GluD1 reduces synaptic GluN2B-containing NMDAR currents, impairing plasticity and cognition. Multiple complementary approaches, including electrophysiological recordings of NMDAR-mediated EPSCs, pharmacological blockade, and immunostaining for surface GluN2B at functional synapses, converge to support the conclusion that GluD1 maintains synaptic GluN2B surface levels and proper NMDAR function. These results extend prior work establishing GluD1 as a trans-synaptic organizer that couples presynaptic neurexin-cerebellin complexes to postsynaptic glutamate receptor function by specifying which NMDAR subtype is maintained at hippocampal synapses (*12, 34*). This advances our understanding of how GluD1 contributes to the molecular architecture underlying synaptic function, synaptic plasticity, and memory.

### The role of GluD1 in controlling GluN2B-containing NMDARs during maturation

NMDARs consist of various subunits, including GluN1, GluN2A-2D, and GluN3A-3B, and it is widely recognized that GluN2A and GluN2B are the primary regulatory subunits of NMDARs (*49*). During early development, GluN2B subunits predominate in NMDARs (*40–42*); however, as development progresses, there is a gradual incorporation of GluN2A subunits in NMDARs, both in the hippocampus and in cultured hippocampal neurons (*40–43, 59*). This is confirmed in our mRNA measurements (Figures 1 and 3). Previous work has shown that mature cortical and hippocampal neurons contain diheteromeric GluN1/GluN2A and GluN1/GluN2B receptors, as well as triheteromeric GluN1/GluN2A/GluN2B receptors, each estimated to comprise approximately one-third of the total NMDAR population (*43, 45*). In our study, after pharmacologically blocking GluN2B-containing NMDARs in control neurons, the remaining responses are primarily mediated by GluN1/GluN2A NMDARs and triheteromeric GluN1/GluN2A/GluN2B NMDARs. The similar NMDAR-EPSC amplitudes and decay kinetics observed in control and GluD1 KO neurons after GluN2B blockade suggest that GluD1 deletion does not substantially alter GluN1/GluN2A-mediated responses, but instead preferentially reduces signaling through GluN2B-containing NMDARs (Figures 1 and 3). Consistent with this interpretation, GluD1 KO neurons showed a selective reduction in GluN2B puncta size, with no corresponding change in GluN2A puncta size (Figure 2). However, because the pharmacological and kinetic approaches used here cannot fully resolve receptor stoichiometry, we cannot fully exclude a contribution from triheteromeric GluN1/GluN2A/GluN2B receptors to the observed effects. An additional limitation is that our measurements do not resolve the contribution of glycine-sensitive GluN3-containing receptors (*71–74*). Recent studies have shown that GluN1/GluN3A excitatory glycine receptors are strongly enriched in the ventral hippocampus (*73*) and that GluN3A deletion accelerates NMDAR decay, suggesting that GluN3A can influence the relative contribution of conventional GluN2A- and GluN2B-containing NMDARs (*75*). However, GluN1/GluN3A receptors are unconventional NMDARs that are activated by glycine and are insensitive to glutamate (*73*). A direct contribution of GluN1/GluN3A receptors to the altered NMDAR-mediated responses observed in GluD1 KO neurons is therefore unlikely, because our evoked NMDAR-EPSCs primarily reflect glutamatergic transmission, whereas NMDA puffing predominantly activates GluN2-containing NMDARs (Figures 1 and 3). Nevertheless, these findings raise the possibility that GluD1 and GluN3A are functionally linked through an as-yet-unknown mechanism that regulates GluN2B-containing NMDARs.

This selective loss of GluN2B-containing NMDARs in GluD1 KO neurons appears to be in contrast with findings from earlier studies of GluD1 function. A previous study reported an increased GluN2B/GluN2A ratio in constitutive GluD1 knockout mice based on bulk biochemical measurements during early postnatal development (*32*). Two non-exclusive factors help reconcile these observations. First, constitutive deletion throughout development likely engages compensatory and homeostatic programs. Constitutive absence of GluD1 can reshape transcriptional profiles and receptor trafficking, potentially increasing GluN2B protein levels or delaying the GluN2B→GluN2A developmental switch at non-synaptic sites. By contrast, our postnatal circuit-specific manipulations minimize long-term compensation and reveal the immediate requirement for GluD1 in maintaining GluN2B at more mature synapses. Second, earlier work quantified total receptor subunits in tissue lysates at P21-P30 (*32*), which pool synaptic and extrasynaptic receptors across heterogeneous cell types and developmental states; our approach isolates functional synaptic NMDARs using electrophysiology and surface-specific immunostaining in defined hippocampal neurons and vCA1→vSub synapses. Together, these differences in timing, readout, and level of analysis suggest a model in which GluD1 participates in GluN2B-NMDAR regulation throughout development, but with distinct roles: during early circuit assembly, its loss may trigger global rebalancing of GluN2B/GluN2A expression, whereas in more mature synapses GluD1 is required cell-autonomously to stabilize synaptic GluN2B-containing NMDARs and preserve GluN2B-NMDAR signaling. This framework predicts that temporally controlled GluD1 deletion across developmental and maturation windows, coupled to subunit-resolved synaptic recordings, will uncover phase-specific functions of GluD1 in sculpting the developmental NMDAR trajectory.

### The role of GluD1 in distinct circuits and inhibitory synapses

GluD1 is broadly expressed in higher brain regions, including hippocampus, cortex, and limbic structures, where it localizes to both excitatory and inhibitory synapses (*8, 62, 63, 76*). Studies in the cerebellum and hippocampus have established that GluD1/2-cerebellin-neurexin (GluD1/2-Cbln-Nrxn) complexes instruct synapse formation and tune inotropic glutamate receptor composition at excitatory synapses (*28, 34, 39, 77–79*). For instance, Tao et al. showed that postsynaptic GluD1 promotes the assembly and maintenance of hippocampal CA3-CA1 excitatory synapses through Cbln2-dependent trans-synaptic interactions (*28*). In contrast, Dai et al. demonstrated that GluD1 can regulate postsynaptic receptor function without altering synapse number in CA1→subiculum synapses. In this pathway, Cbln2–Nrxn3 and Cbln2–Nrxn1 complexes differentially regulate AMPAR- and NMDAR-mediated transmission through distinct postsynaptic GluD1 signaling mechanisms (*34, 80*). Our present data further define a pathway-specific excitatory function of GluD1 at vCA1→vSub synapses, where it is required to maintain the synaptic GluN2B-containing NMDAR component and GluN2B-dependent LTP. Together with reports that GluD1 knockdown can spare LTP at CA3-CA1 synapses (*28*), these findings support a circuit-specific model in which distinct GluD1-containing trans-synaptic complexes confer unique rules for receptor-subtype maintenance and plasticity. Such specificity may arise from differences in cerebellin isoforms, neurexin family members, or neurexin splice variants across synaptic pathways. Because neurexin-ligand combinations generate diverse synaptic properties (*12*), GluD1-dependent stabilization of GluN2B-containing NMDARs may represent one pathway-specific component of a broader synaptic-organizer program.

In parallel, GluD1 has emerged as an important organizer of inhibitory synapses. GluD1 is required for the formation and function of specific dendritic GABAergic inhibitory synapses, regulates inhibitory transmission, and supports inhibitory plasticity (*27, 76, 81*). Fossati et al. demonstrated that postsynaptic GluD1 interacts with Cbln4 released from somatostatin-positive interneurons and with presynaptic neurexins to promote a defined population of dendritic inhibitory synapses (*81*). More recently, Piot et al. showed that GluD1 localizes to hippocampal GABAergic synapses, directly binds GABA through its ligand-binding domain, and mediates a long-lasting, non-ionotropic potentiation of inhibitory transmission that depends on trans-synaptic anchoring (*27*). Thus, the synaptic function of GluD1 is likely determined by circuit context, synapse type, and the specific complement of cerebellin and neurexin partners.

The present electrophysiological and imaging experiments specifically interrogated excitatory NMDAR-mediated transmission and therefore provide direct evidence for an excitatory postsynaptic function of GluD1 at vCA1→vSub synapses. However, these experiments were not designed to exclude parallel effects of GluD1 deletion at inhibitory synapses. Disruption of GluD1-dependent GABAergic synapse organization or inhibitory plasticity could alter dendritic integration, excitation–inhibition balance, NMDAR activation thresholds, and the induction or expression of plasticity in vSub neurons. Such changes could also contribute to the broader circuit-level and behavioral phenotypes observed *in vivo*. We therefore interpret the reduced NMDAR responses and impaired LTP as primarily resulting from disrupted excitatory GluD1 signaling at vCA1→vSub synapses. However, the broader physiological and behavioral effects of GluD1 deletion may reflect combined alterations in excitatory GluN2B-containing NMDAR function and inhibitory synaptic regulation. Future studies using synapse-type-, cell-type-, and interneuron-specific manipulations will be required to determine the relative contributions of these excitatory and inhibitory GluD1 functions within the ventral hippocampal circuit.

### Potential mechanisms by which GluD1 controls GluN2B-containing NMDARs

Mechanistically, our findings support a model in which GluD1 regulates the synaptic retention and/or trafficking of GluN2B-containing NMDARs rather than simply modulating global expression. Prior structural and functional work has shown that GluD1 operates as a signal-transduction hub: transsynaptic neurexin-cerebellin complexes engage its large extracellular domain, while its intracellular C-terminal tail (CT) connects to unknown postsynaptic scaffolds and signaling machinery (*14, 15, 34, 82*). Notably, the neurexin-1 and cerebellin-2 controls the NMDAR-mediated LTP (*38, 39*), providing a plausible transsynaptic mechanism through which GluD1 could influence GluN2B-dependent Ca²⁺/CaMKII signaling and LTP (Figure S5). At the postsynaptic site, mutations in GluD1-CT motifs alter synaptic responses in a manner that phenocopies GluD1 loss (*34*), suggesting that CT-bound effectors are required to propagate GluD1-dependent signals.

One plausible mechanism is that GluD1-CT-dependent signaling facilitates the capture or retention of GluN2B-containing NMDARs within postsynaptic scaffolding complexes. These scaffolds regulate NMDAR transport, surface distribution, lateral mobility, and stabilization at postsynaptic sites (*46, 83–85*). GluD1 could therefore stabilize GluN2B-containing NMDARs indirectly by recruiting an unidentified CT-binding protein that connects GluD1 signaling to scaffold complexes. A second, nonexclusive mechanism is altered phosphorylation-dependent trafficking. For instance, Src-family kinases promote GluN2B surface expression and synaptic stabilization (*86–89*). GluD1 loss could disrupt this pathway, enhance receptor internalization, and reduce synaptic GluN2B availability without changing total receptor abundance. A third possibility is that GluD1 regulates GluN2B-containing NMDARs through metabotropic signaling. GluD1 has been linked to mGlu1/5-dependent signaling at excitatory synapses, and disease-associated GRID1 variants disrupt this pathway and impair synaptic function (*20, 22, 33, 37, 90*). mGluRs can regulate NMDAR phosphorylation, trafficking, and synaptic responses (*91–93*). GluD1 could therefore influence GluN2B trafficking indirectly by organizing a local mGlu1/5-dependent signaling complex. Finally, GluD1-dependent signaling could potentially influence the gating or allosteric state of GluN2B-containing NMDARs, because intracellular signaling and GluN2B C-terminal modifications are known to regulate NMDAR channel behavior (*94–96*). However, this possibility appears less likely because our data primarily demonstrate an overall reduction in surface GluN2B-containing NMDARs rather than changes in the functional properties of individual receptors.

Together, we propose a working model in which presynaptic neurexin–cerebellin complexes engage postsynaptic GluD1, whose C-terminal tail coordinates scaffolding, phosphorylation, and receptor trafficking pathways to favor the synaptic retention and stabilization of GluN2B-containing NMDARs at mature excitatory synapses (Figure S5). Identifying GluD1-CT interactors will be an important next step to pinpoint the molecular machinery mediating subtype-specific control.

### The role of GluD1 in cognitive processes and disease

The strong impact of GluD1 on GluN2B-containing NMDAR-dependent plasticity and contextual fear memory in our study resonates with converging human and animal data implicating GRID1 in cognition and neuropsychiatric disease (*20, 23*). In addition to memory deficits, GluD1 manipulations have been associated with altered emotional behaviors, including anxiety/depression-like phenotypes (*23, 97–99*), consistent with the prominent role of ventral subicular circuits in regulating anxiety-related responses (*64, 100, 101*). Thus, our findings that GluD1 controls GluN2B-containing NMDAR function at vCA1→vSub synapses suggest a circuit-level mechanism by which GRID1 dysfunction could jointly impact cognitive and anxiety-related dimensions of disease.

Our work adds mechanistic depth to these associations by identifying a GluD1-dependent synaptic pathway in which loss of GluD1 is accompanied by reduced GluN2B-containing NMDAR signaling, impaired synaptic plasticity, and deficits in long-term contextual memory. However, the present data do not establish that the behavioral phenotype is mediated solely by reduced GluN2B-containing NMDAR function. Although learning ability remains preserved, suggesting that task acquisition can be supported by parallel or compensatory circuits, GluD1 signaling in the vCA1→vSub pathway may be particularly important for the stabilization and retrieval of learned information. Future experiments using acute local suppression of GluN2B-containing NMDAR signaling in the ventral subiculum will be important to determine whether this manipulation phenocopies aspects of the GluD1 deletion phenotype and more directly links receptor dysfunction to behavior.

NMDAR subunit composition is a well-established determinant of synaptic plasticity and cognitive functioning, and pathogenic variants in NMDAR subunit genes cause severe neuropsychiatric disorders with intellectual disability and schizophrenia (*46, 48, 102–104*). By showing that GluD1 controls GluN2B-containing NMDARs specifically, our data suggest that GluD1-related conditions may represent an “organizer-level” NMDARopathy, in which signaling and plasticity are perturbed because receptor complexes are mis-positioned or mis-specified, rather than intrinsically mutated. This conceptual shift broadens the therapeutic landscape: instead of only targeting NMDARs directly, which can be limited by side effects, one could attempt to restore GluD1-dependent organizer pathways to normalize GluN2B-containing NMDAR function in vulnerable circuits. In summary, clarifying the GluD1-NMDAR axis will fill a key mechanistic gap between synaptic architecture and plasticity, address synaptic pathology underlying GluD1-linked cognitive and anxiety-related deficits, and highlight tractable targets for therapeutic intervention.

## METHODS

### Mice

All mice were maintained on a C57BL/6 background under protocols approved by the Mount Sinai Icahn School of Medicine Institutional Animal Care and Use Committee. Animals were group-housed with same-sex littermates on a 12-hr light/dark cycle with ad libitum access to food and water. CRISPR/Cas9 knockin mice (Jackson Laboratory, Jax Stock No. 026179) were genotyped by PCR using IDT primers: for Cas9KI, forward 5′-GCTAACCATGTTCATGCCTTC-3′ and reverse 5′-CTCCGTCGTGGT CCTTATAGT-3′; for Cas9WT, forward 5′-CTGGCTTCTGAGGACCG-3′ and reverse 5′-AGCCTGCCCAGAAGACTCC-3′. For synaptic functional studies, both male and female littermates were used; behavioral studies employed male mice. Littermates were always used as control. All data collection and analyses were performed under blinded conditions, except for the *in vivo* CRISPR-mediated deletion of GluD1 in the subiculum (Figure 3), for which mCherry/tdTomato expression was required to distinguish control from infected neurons, and for mRNA (Figures 1A and 3A) and immunoblotting (Figures 2A and S1) measurements, which provided objective quantitative readouts independent of the observer.

### DNA constructs and Viruses

Lentiviral or adeno-associated viruses (AAVs) were used in cultured neurons and *in vivo* subiculum injections as described previously (*34*). Briefly, CRISPR/Cas9-mediated GluD1 knockout was achieved with a validated Lenti-sgRNA following Dai et al., 2021 (*34*) (Figure S1) or AAV-DJ construct expressing either control guides under U6 and H1 promoters alongside an hSyn-driven reporter, following Wang et al., 2021 (*105*) (Figure 3A). For *in vitro* culture experiments, lentiviral vectors bearing the same guide cassettes were produced in HEK293T cells (ATCC) via calcium-phosphate method: each 75 cm² plate was transfected with 12 µg of the lentiviral expression plasmid (control or Grid1-targeting guide) plus 12 µg each of pRSV-REV, pMDLg/pRRE, and pVSVG. After overnight, cells were washed with warm PBS and switched to neuronal growth medium; 48 hours post-transfection, supernatant was collected, centrifuged at 1,500g for 10 minutes to remove debris, filtered through a 0.22 µm membrane, aliquoted, and stored at −80 °C until use. For sparse labeling *in vivo*, we further concentrated our lentivirus as descried before (*34*). High-titer AAV stocks were produced by the University of North Carolina Vector Core and virus preparations were diluted to the appropriate working concentration immediately prior to injection to achieve the desired expression levels.

### Hippocampal Cultures

Hippocampal cultures were prepared from randomly selected P0 pups as described previously (*34, 38*): hippocampi were dissected in ice-cold HBSS, irrespective of sex, digested in 10 U·mL⁻¹ papain (Worthington) at 37 °C for 20-25 min, then rinsed with prewarmed plating medium (MEM with 2 mM L-glutamine, 0.4% D-glucose, 2% B-27® Supplement, and 5% FBS). Tissue was triturated in plating medium, and neurons were seeded onto Matrigel-coated coverslips in 24-well plates (six hippocampi per one plate), designating this as DIV0. On DIV1, 95% of plating medium was replaced with prewarmed neuronal growth medium (Neurobasal-A with 2 mM L-glutamine, 2% B-27® Supplement, and 5% FBS). At DIV3, half of the medium was exchanged with growth medium containing 4 µM AraC to limit glial proliferation. On DIV4, cultures were randomly assigned and transduced with either control or *Grid1*-targeting AAVs or lentiviruses. All functional and morphological analyses were conducted at DIV15-17.

### mRNA measurements

mRNA was prepared from hippocampal cultures harvested at DIV1,4,9,16, or brain tissue from the hippocampus region of P0,7,21,60 mice as previously described (*34, 38*). RNA extraction was performed using TRIzol (Thermo Fisher, 15596026) and quantified using an ND-1000 spectrophotometer (NanoDrop, ThermoScientific). Quantitative real-time PCR (RT-qPCR) was performed using the TaqMan Fast Virus 1-Step Master Mix (Applied Biosystems) based on the manufacturer’s instructions, and reactions were carried out and quantified using a QuantStudio 7 Pro instrument (Applied Biosystems). Expression levels were normalized to *Actb* as an internal control. The following PrimeTime qPCR Assays (IDT) were used (shown as gene, primer1, probe, and primer2 or predesigned): *Grid1*, TGGATATGCCAGTGCGTTAC, TCCATCATCCACATCGGTGCCATC, CAGGTCTGATACAGCCAACTG; *Grin1* (Mm.PT.58.330045830); *Grin2a* (Mm.PT.58.32291523); *Grin2b* (Mm.PT.58.42676841); *Actb* (Mm.PT.58.33540333).

### Cell line identification

HEK 293T cells were directly purchased from ATCC, which regularly validates cell lines. Cell lines were tested negative for mycoplasma contamination using the fluorochrome Hoechst DNA stain and the direct culture method (*106–108*).

### Immunoblotting

The protocol was conducted as previously described (*34, 38*). Neuronal cultures were lysed for immunoblotting by aspirating medium, washing once with prewarmed PBS, and adding Laemmli buffer (12.5 mM Tris-HCl pH 6.8, 5 mM EDTA pH 6.8, 143 mM β-mercaptoethanol, 1% SDS, 0.01% bromophenol blue, 10% glycerol), after which samples were boiled, separated by SDS-PAGE at 120 V for ∼1.5 hours, and transferred to nitrocellulose membranes via the Trans-Blot Turbo system (Bio-Rad). Membranes were blocked in 5% milk in TBS with 0.1% Tween-20 (TBST) at room temperature for 1 hour, incubated overnight at 4 °C with primary antibodies (anti-GluD1 rabbit (1:1,000; Frontier Institute; Cat#Af1390); anti-GluN1 mouse, 1:500, Synaptic Systems Cat#114011; anti-GluN2A mouse, 1:500, Invitrogen Cat#MA5-27692; anti-GluN2B rabbit, 1:500, Invitrogen Cat#71-8600; anti-Tuj1 rabbit, 1:2,000, Sigma-Aldrich Cat#T2200), washed three times with TBST, then incubated with IR-dye-conjugated donkey secondary antibodies (anti-rabbit, anti-mouse, anti-guinea pig, each 1:10,000; LI-COR Biosciences). Blots were scanned on an Odyssey Infrared Imager and quantified using Odyssey software, with protein intensities normalized first to Tuj1 and then to the relevant control samples.

### Immunocytochemistry

The protocol was conducted as previously described (*34, 38*). Hippocampal cultures were fixed in 4% paraformaldehyde with 4% sucrose for 10 min, washed three times with PBS, and blocked in 5% goat serum in PBS for 1 h at room temperature; for surface labeling, coverslips were incubated overnight at 4 °C with N-terminal antibodies diluted in blocking buffer (anti-GluN1, 1:300, mouse, Synaptic Systems Cat#114011; anti-GluN2A, 1:300, mouse, Invitrogen Cat#MA5-27692; anti-GluN2B, 1:300, rabbit, Invitrogen Cat#71-8600), then washed three times with PBS and permeabilized in 5% goat serum with 0.3% Triton X-100 for 1 h at room temperature. Following permeabilization, coverslips were incubated overnight at 4 °C with intracellular markers diluted in blocking buffer (anti-vGluT1, 1:1000, guinea pig, Millipore Cat# AB5905; anti-PSD95, 1:1000, mouse, Invitrogen Cat#MA1-046 or rabbit, Alomone Cat#APZ-009; anti-MAP2, 1:1000, chicken, Millipore Cat#AB15452), washed three times in PBS, and incubated for 1 h at room temperature with species-appropriate Alexa Fluor secondary antibodies (1:1000 for Alexa Fluor 405, 488, 546, 647). After three PBS washes, coverslips were mounted in Fluoromount-G (SouthernBiotech). For image acquisition, three or more independent cultures were used with at least three coverslips in total per condition; confocal images were collected at room temperature on a Zeiss LSM 880 with a 63× oil-immersion objective at 2048 × 2048 pixel resolution, acquiring 13 optical sections at 0.25 µm z-steps and generating maximum-intensity projections. Images were converted using ImageJ/Fiji (NIH) (*109*) to Nikon Images file (.nd2), then analyzed using NIS-Elements AR Analysis software (Nikon). Synaptic puncta were quantified by thresholding to exclude background and selecting puncta in the size range 0.04-3 µm² to analyze.

### Stereotactic Injections

Stereotactic injections of AAV or lentivirus into mice at P21-23 were performed essentially as described (*34, 38, 69, 110*). Mice were anesthetized with ketamine (90 mg/kg) and xylazine (10 mg/kg) and placed in a stereotaxic frame (Harvard Apparatus). Using a Hamilton syringe, 0.9 μL of AAVs (∼10^8-10^9 GC/mL) or concentrated lentivirus was delivered at a rate of 0.1 μL/min into the ventral subiculum (coordinates relative to Bregma: AP −3.4 mm, ML ±3.2 mm), with 0.9 μL dispensed at two dorsoventral depths (DV −3.45, and −3.2 mm). Littermates were randomly assigned to receive either control or sgRNA-expressing virus. After injection, the needle was left in place for ∼5 min to minimize backflow before slow withdrawal. Viral expression was confirmed by mcherry or tdTomato fluorescence: example images were acquired on a Zeiss LSM 880 confocal microscope with a 10× objective at 1024 × 1024 resolution (Figure S3A), and all slices used for electrophysiology were screened under an Olympus fluorescence microscope.

### Culture Electrophysiology

Electrophysiological recordings were conducted on hippocampal neurons cultured from Cas9 P0 mice, infected at DIV4-5 with either Lenti-sgRNA-mCherry or AAV-sgRNA-tdTomato for GluD1 knockout (with Lenti-mCherry or AAV-tdTomato as controls) and analyzed at DIV15-17 as previous described (*34*). During recording, cultures were superfused at room temperature with ACSF (in mM: 120 NaCl, 2.5 KCl, 1 NaH₂PO₄, 26.2 NaHCO₃, 2.5 CaCl₂, 1.3 MgSO₄·7H₂O, 11 d-glucose; ∼290 mOsm), continuously bubbled with 95% O₂/5% CO₂. To isolate NMDAR-mediated currents, ACSF contained 50 µM picrotoxin and 10 µM CNQX, and cells were held at 0, +20, +40, and +60 mV. To preferentially inhibit diheteromeric GluN1/GluN2B receptors while minimizing effects on triheteromeric GluN1/GluN2A/GluN2B receptors, we used a low concentration (1 µM) of the GluN2B-selective antagonist Ro 25-6981 (*53–55*). Patch pipettes were filled with internal solution (in mM: 117 Cs-methanesulfonate, 15 CsCl, 8 NaCl, 10 TEA-Cl, 0.2 EGTA, 4 Na₂-ATP, 0.3 Na₂-GTP, 10 HEPES; pH 7.3 adjusted with CsOH; ∼300 mOsm). Synaptic responses were evoked by a nichrome stimulating electrode placed 100-150 µm from the soma, controlled by a Model 2100 Isolated Pulse Stimulator synchronized with Clampex 10, and recorded using a MultiClamp 700B amplifier with signals digitized at 4 kHz.

### Slice Electrophysiology

Electrophysiological recordings in acute hippocampal slices were performed 6-9 weeks after stereotactic lentivirus or AAV injection into the subiculum of Cas9 mice as previously described (*34, 38, 39*). Horizontal hippocampal slices (300 µm) were prepared in ice-cold, high-sucrose cutting solution (in mM: 85 NaCl, 75 sucrose, 2.5 KCl, 1.3 NaH₂PO₄, 24 NaHCO₃, 0.5 CaCl₂, 4 MgCl₂, 25 D-glucose), then equilibrated in ACSF at 31 °C for 30 min followed by 1 h at room temperature. During recordings, slices were superfused at room temperature with ACSF (in mM: 120 NaCl, 2.5 KCl, 1 NaH₂PO₄, 26.2 NaHCO₃, 2.5 CaCl₂, 1.3 MgSO₄·7H₂O, 11 D-glucose; ∼290 mOsm) bubbled with 95% O₂/5% CO₂. Synaptic responses in subiculum neurons were evoked by placing a nichrome stimulating electrode at the distal CA1 region. For NMDAR current-voltage (I-V) curves recordings, cells were voltage-clamped using an internal solution (in mM: 117 Cs-methanesulfonate, 15 CsCl, 8 NaCl, 10 TEA-Cl, 0.2 EGTA, 4 Na₂-ATP, 0.3 Na₂-GTP, 10 HEPES; pH 7.3 adjusted with CsOH; ∼300 mOsm). All NMDAR-EPSCs were recorded in the presence of 50 µM picrotoxin and 10 µM CNQX, and cells were held at 0, +20, +40, and +60 mV. To preferentially inhibit diheteromeric GluN1/GluN2B receptors while minimizing effects on triheteromeric GluN1/GluN2A/GluN2B receptors, we used a low concentration (1 µM) of the GluN2B-selective antagonist Ro 25-6981 (*53–55*). For LTP recordings, cells were voltage-clamped using either the same internal as for NMDAR I-V curves or with an internal solution (in mM: 137 K-gluconate, 5 KCl, 10 HEPES, 4 ATP-Mg_2_, 0.5 GTP-Na_2_, 10 phosphocreatine, 0.2 EGTA, pH 7.2 with KOH; ∼300 mOsm). Neuron firing type (burst-vs. regular-spiking) was determined in current-clamp immediately after break-in by injecting depolarizing current. All LTP were done in the presence of 50 µM picrotoxin with holding potential at −70 mV. LTP was induced under voltage clamp at 0 mV by four 100 Hz, 1 s stimulus trains with 10 s inter-train intervals; baseline (last 5 min pre-LTP) and post-LTP (last 5 min) responses were recorded at 0.1 Hz. Paired-pulse ratios (40 ms interval) were measured before and after LTP induction. For NMDA puffing experiment, application of NMDA (Tocris Bioscience, in the presence of picrotoxin and CNQX) was performed with 10 psi for 10-200 ms by using Picospritzer III (Parker Instrumentation), and cells were held at +40 mV. The total charge was calculated within 25 s from puff application. The slope was calculated from data obtained between 10 ms to 200 ms. Data were analyzed using Clampfit and Igor Pro.

### Open Field Test

The protocol was conducted as previously described (*111, 112*). Animals were placed in a 44 × 44 × 44 cm square chamber constructed from closed-cell foam with a sealed white surface for easy cleaning and optimal video contrast. Each subject was allowed to freely explore the arena for 10 minutes under red light. All chambers were thoroughly hand-cleaned between tests with a Peroxigard 0.5% hydrogen peroxide solution. The floor was virtually divided into a central zone (22 × 22 cm) and four corner zones (11 × 11 cm each). Measured parameters included time spent in the center, number of entries into the center zone, and total distance. All tracking and scoring were performed using Viewer 3 software (Bioserve).

### Elevated Plus Maze

The protocol was conducted as previously described (*111, 112*). Animals were placed in an elevated plus maze (EPM) consisting of a white acrylic plus-shaped platform with two open arms (30 × 5 cm) and two closed arms of the same dimensions with 15 cm-high white walls, elevated 50 cm above the floor. Each subject was allowed to freely explore the maze for 10 minutes under red light. EPM was thoroughly hand-cleaned between tests with a Peroxigard 0.5% hydrogen peroxide solution. Measured parameters included total time spent in the open arms and total time spent in the closed arms. All tracking and scoring were performed using Viewer 3 software (Bioserve).

### Three Chamber Sociability Test

The protocol was conducted as previously described (*113, 114*). A three-chamber arena of white acrylic (approximately 60 × 40 × 22 cm total, with three equal ∼20 × 40 × 22 cm compartments) is separated by removable clear acrylic doors (∼5 × 5 cm openings), and each lateral chamber holds a ventilated container (∼10 cm diameter × 15 cm height) with bars for limited interaction. All tests were under red light. Mice are habituated one day before by freely exploring all chambers for 5 minutes. On the test day, after cleaning, each subject underwent three consecutive 5-minute tests: during test0, the subject mouse was placed the mouse in the center with identical empty containers in both lateral chambers and allowed to freely explore the chambers (to assess side bias); during test1, a juvenile (P21-28) stimulus mouse was presented in one container, while the opposite container remained empty; during test2, a second novel juvenile stimulus mouse was presented in the previously empty chamber while the first (now familiar) juvenile stimulus mouse remains. All chambers were thoroughly hand-cleaned between tests with a Peroxigard 0.5% hydrogen peroxide solution. Using Viewer 3 (Bioserve), time spent in each lateral chamber during each test was recorded, and social index (SI) ratios were computed by first normalizing each chamber’s time to its corresponding prior trial to account for any inherent chamber preference, then comparing time spent with the social versus non-social target within each test. The original calculation of the discrimination index is indicated in Figure S4 (*113, 115*).

### Two-Chamber Avoidance Test

The modified protocol was conducted as previously described (*38, 39*) using a Shuttle Box (Med Associates, Inc.) comprising two visually distinct chambers under dim lighting separated by a gate. Habituated one day prior, mice then enter training day (as described below). The right chamber delivered a foot shock (0.25 mA, 2 s) after a 2-second delay upon entry, prompting immediate retreat to the left chamber; mice were placed initially in the left chamber and allowed to explore freely, with each entry into the right chamber and its latency recorded as a trial. Training continued until the mouse remained in the left chamber for over 2 minutes without re-entering the shock-paired side. Contextual memory test was assessed at 1 and 7 days by reintroducing the mouse to the left chamber for a 2-minute test period, during which entries into—and latency to enter—the right chamber were measured.

### Data Analysis

Statistical significance was determined using Student’s t-test or one-way or two-way analysis of variance (ANOVA), followed by Tukey’s post hoc multiple-comparisons test, as appropriate. In cases where one group within a comparison did not meet the normality assumption, the corresponding analysis was also performed using the nonparametric Mann–Whitney U test or Kruskal–Wallis test followed by Dunn’s multiple-comparisons test. Statistical significance is indicated as * = *p*<0.05, ** = *p*<0.01 and *** = *p*<0.001. Nonsignificant results (*p* > 0.05) are not specifically indicated.

## ACKNNOWLEGEMENTS

This work was supported by the Brain & Behavior Research Foundation Grant NARSAD31414 (to J.D.), Alkermes Award 74965 (to J.D.), National Institutes of Health Grant R01HL167520 (to E.S.), T32MH087004 (to D.L.S.), T32GM154814 (to C.V.), and ISMMS faculty start up fund (to J.D.). This work was also supported by the Marie Skłodowska-Curie Action Innovative Training Network ‘Serotonin and Beyond’ under the European Union’s Horizon 2020 research and innovation program (grant agreement 953327 to M.W.) and the Fonds Wetenschappelijk Onderzoek - Vlaanderen (FWO; Research Foundation - Flanders; grant agreement G0E66722N to M.W.). We thank the Friedman Brain Institute and the Departments of Pharmacological Sciences and Neuroscience at the Icahn School of Medicine at Mount Sinai for their support. We appreciate members of the Dai lab for experimental assistance and helpful discussions. We thank Drs. William Janssen and Mustafa Siddiq (ISMMS Microscopy Core) for training and assisting with confocal imaging. We thank Drs. Thomas C. Südhof (Stanford University) and Christopher Patzke (University of Notre Dame) for contributing plasmids. We thank Drs. Nan Yang (ISMMS) and Daniel Wacker (ISMMS) for contributing HEK293T cells.

## AUTHOR CONTRIBUTIONS

E.P. performed all experiments except for the electrophysiological recordings, which were carried out by D.L.S., J.S., T.W., and J.D. M.Z. organized the confocal image analysis. C.V. assisted with mouse line maintenance and stereotaxic injections. E.P., D.L.S., and J.D. designed the experiments and analyzed the data. E.P. and J.D. assembled the figures and wrote the manuscript with input from D.L.S., J.S., T.W., E.S., and M.W.

## Materials and Methods

### KEY RESOURCES TABLE

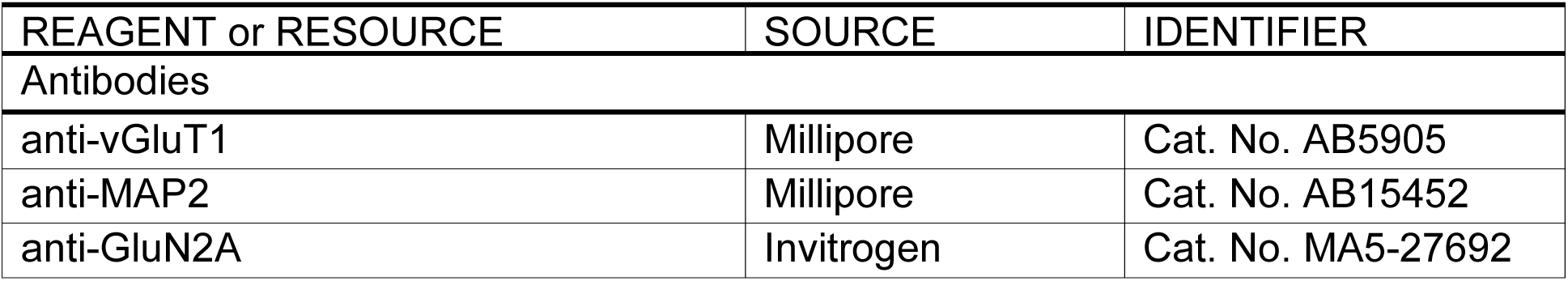
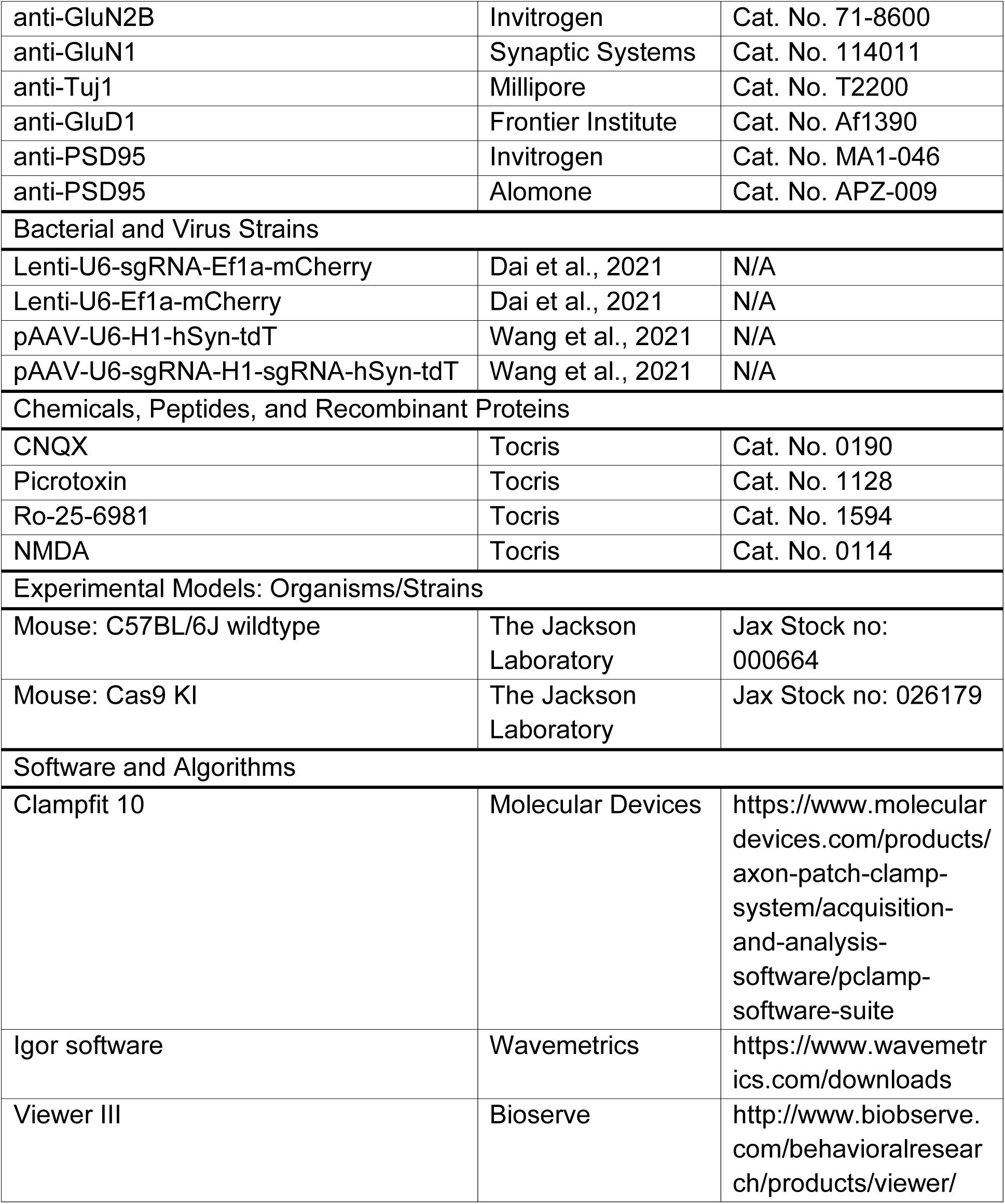
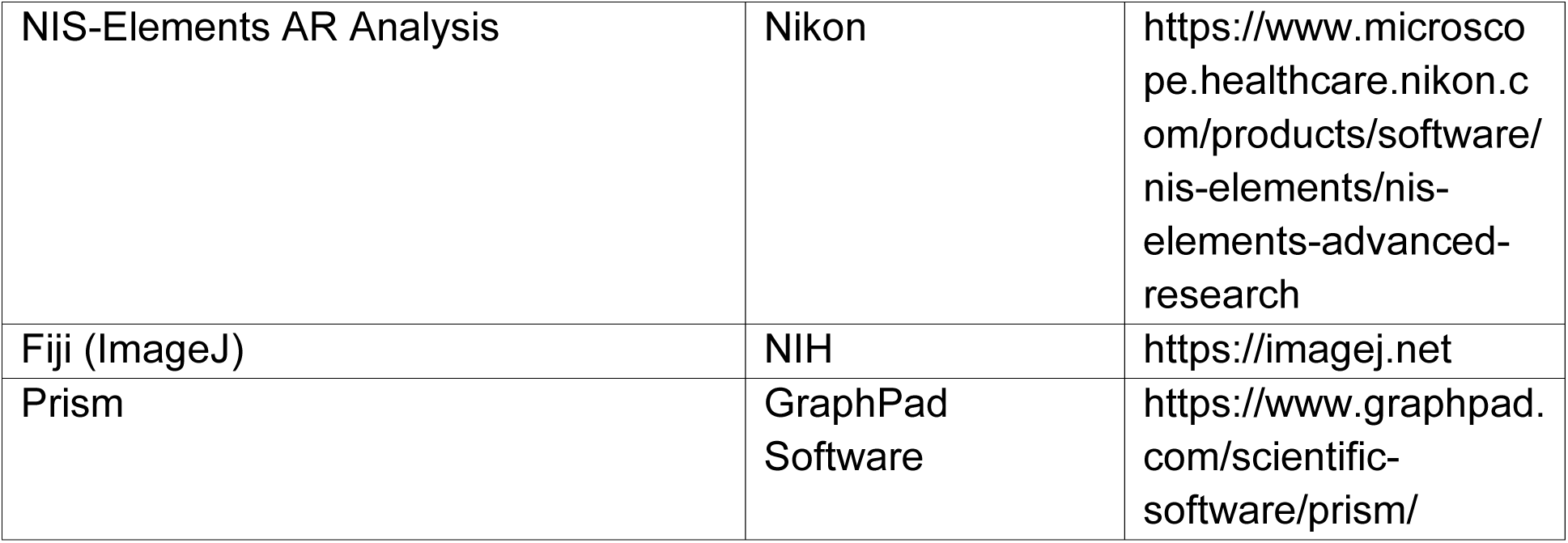

| REAGENT or RESOURCE | SOURCE | IDENTIFIER |
| --- | --- | --- |
| Antibodies |  |  |
| anti-vGluT1 | Millipore | Cat. No. AB5905 |
| anti-MAP2 | Millipore | Cat. No. AB15452 |
| anti-GluN2A | Invitrogen | Cat. No. MA5-27692 |
| anti-GluN2B | Invitrogen | Cat. No. 71-8600 |
| anti-GluN1 | Synaptic Systems | Cat. No. 114011 |
| anti-Tuj1 | Millipore | Cat. No. T2200 |
| anti-GluD1 | Frontier Institute | Cat. No. Af1390 |
| anti-PSD95 | Invitrogen | Cat. No. MA1-046 |
| anti-PSD95 | Alomone | Cat. No. APZ-009 |
| Bacterial and Virus Strains |  |  |
| Lenti-U6-sgRNA-Ef1a-mCherry | Dai et al., 2021 | N/A |
| Lenti-U6-Ef1a-mCherry | Dai et al., 2021 | N/A |
| pAAV-U6-H1-hSyn-tdT | Wang et al., 2021 | N/A |
| pAAV-U6-sgRNA-H1-sgRNA-hSyn-tdT | Wang et al., 2021 | N/A |
| Chemicals, Peptides, and Recombinant Proteins |  |  |
| CNQX | Tocris | Cat. No. 0190 |
| Picrotoxin | Tocris | Cat. No. 1128 |
| Ro-25-6981 | Tocris | Cat. No. 1594 |
| NMDA | Tocris | Cat. No. 0114 |
| Experimental Models: Organisms/Strains |  |  |
| Mouse: C57BL/6J wildtype | The Jackson Laboratory | Jax Stock no: 000664 |
| Mouse: Cas9 KI | The Jackson Laboratory | Jax Stock no: 026179 |
| Software and Algorithms |  |  |
| Clampfit 10 | Molecular Devices | https://www.moleculardevices.com/products/axon-patch-clamp-system/acquisition-and-analysis-software/pclamp-software-suite |
| Igor software | Wavemetrics | https://www.wavemetrics.com/downloads |
| Viewer III | Bioserve | http://www.biobserve.com/behavioralresearch/products/viewer/ |
| NIS-Elements AR Analysis | Nikon | https://www.microscope.healthcare.nikon.com/products/software/nis-elements/nis-elements-advanced-research |
| Fiji (ImageJ) | NIH | https://imagej.net |
| Prism | GraphPad Software | https://www.graphpad.com/scientific-software/prism/ |

