## Supplemental Figures for "GluD1 Modulates GluN2B-containing NMDAR Function and Plasticity at Subicular Synapses"

### SUPPLEMENTARY FIGURES and FIGURE LEGENDS

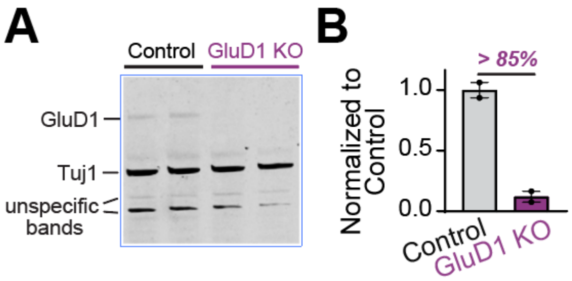


**Figure S1. CRISPR-mediated GluD1 KO in Hippocampal Cultured Neurons of Cas9 Mice at DIV17 Using Lentivirus Infection**

(A and B) Representative immunostaining blot showing protein expression levels of GluD1 and Tuj1 from DIV17 cultured hippocampal neurons. We utilized Cas9 neurons with lentivirus infections of CRISPR sgRNA to delete GluD1. Quantification of relative protein expression levels in GluD1 condition (n = 2, normalized to control conditions after normalized to Tuj1) compared to the control condition (n = 2). Data are mean ± s.e.m.

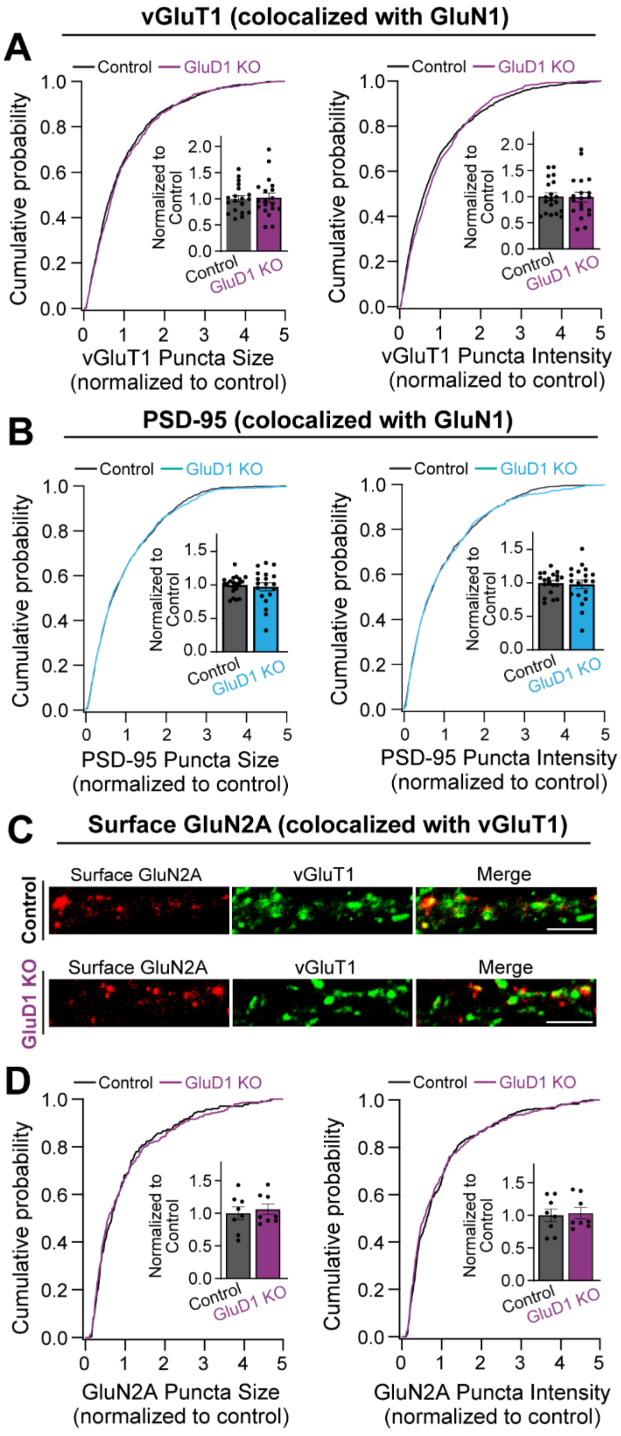


**Figure S2. GluD1 KO does not Alter vGluT1 or PSD-95 Puncta Size and Intensity, also does not Change GluN2A Subunit Surface Expression**

(A and B) Summary graphs and cumulative probability graphs of normalized puncta size and intensity for vGluT1 (A, n = 19 per condition) or PSD-95 (B, n = 20 control and n = 19 KO) from 3 batches of coverslips per condition.

(C and D) Synaptic surface staining and confocal imaging of GluN2A subunit in cultured hippocampal neurons. Representative images of dendrites in Control and GluD1 KO conditions. bar: 5 μm. Summary graphs and cumulative probability graphs of normalized puncta size and intensity for GluN2A (n = 8 per condition) from 3 coverslips per condition.

Unpaired two-tailed t-test or Mann-Whitney U test shows no significant difference. Kolmogorov-Smirnov test was performed for cumulative probability graphs. Data are mean ± s.e.m.

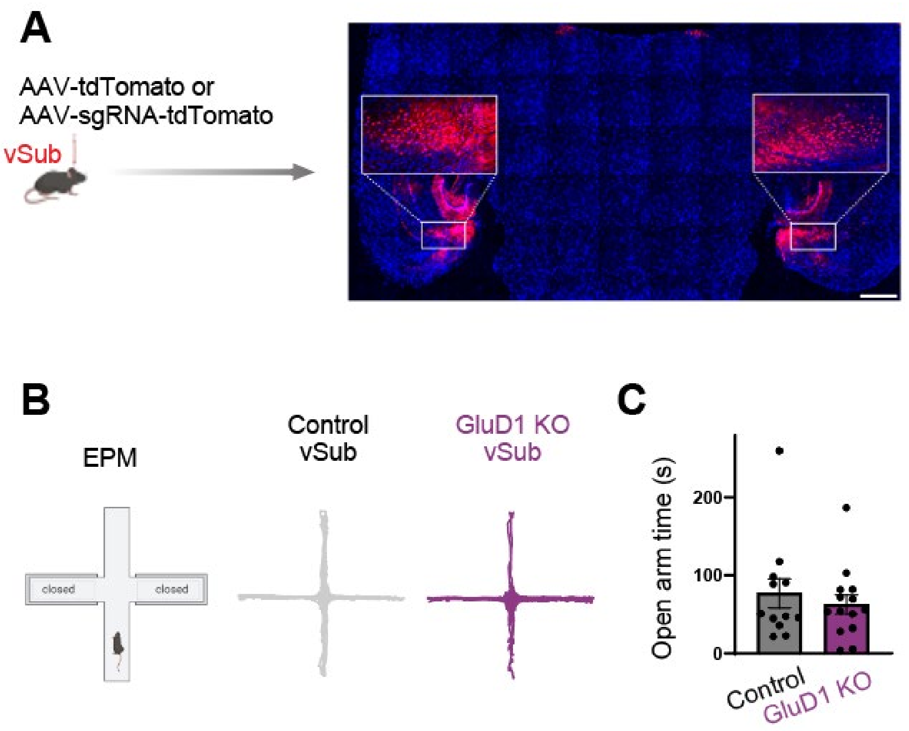


**Figure S3. AAVs Infection in vSub and CRISPR-mediated GluD1 KO in Hippocampal vSub does not Impact Open Arm Time in the EPM Test**

(A) Left: diagram of stereotactic injections. Right: Hippocampal subiculum regions of Cas9 mice were bilaterally infected by stereotactic injections of AAVs expressing tdTomato or sgRNA-tdTomato at P21-23, and subiculum synapses were analyzed at 6-9 weeks. Bar: 500 $\mu$m.

(B and C) Representative EPM traces of Control and GluD1 KO conditions. Summary graphs of open-arm time during the 10-min test. We also analyzed the first 5 min, which yielded similar results (plots not shown). Unpaired two-tailed Mann-Whitney U test shows no significant difference. Data are mean ± s.e.m.

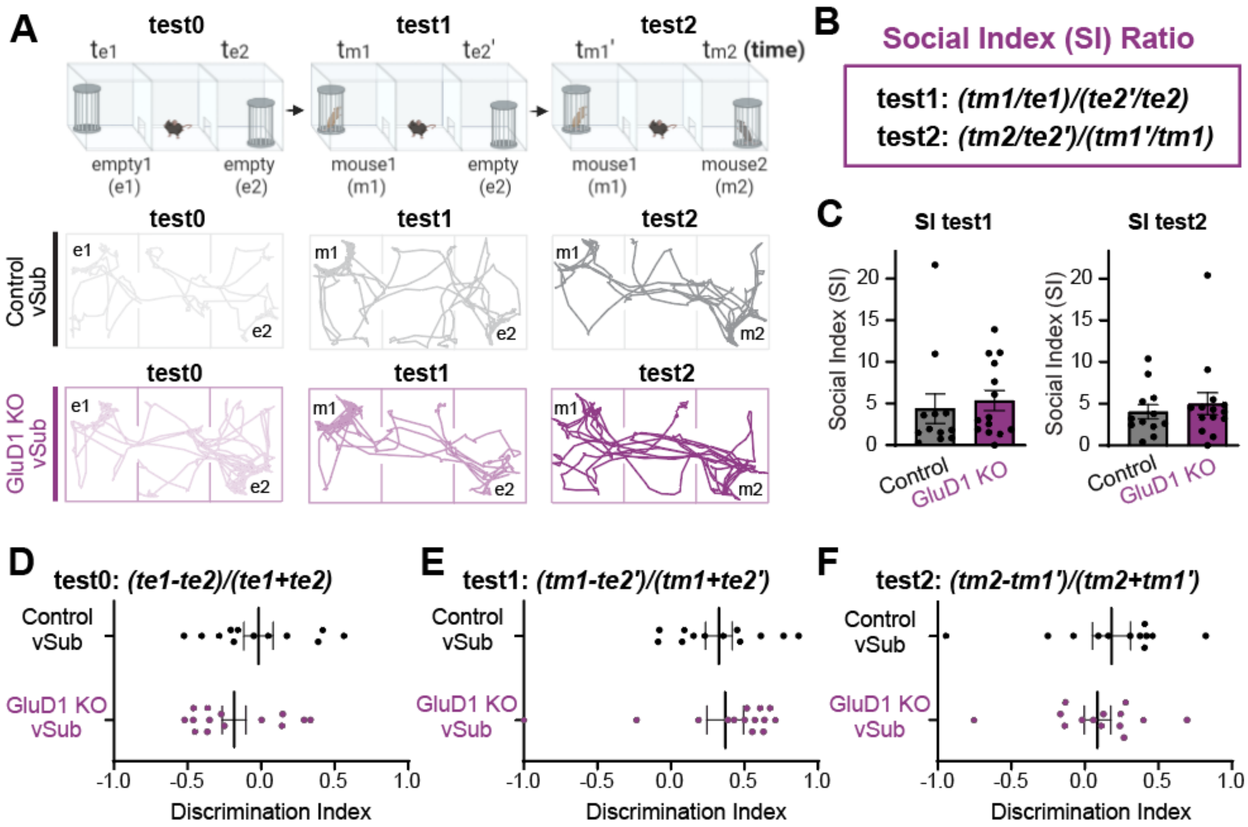


**Figure S4. CRISPR-mediated GluD1 KO in Hippocampal vSub does not Impact Social Behavior in the 3-Chamber Test**

(A) Schematic of the 3-chamber test. Test0: two empty cages; Test1: a stranger mouse in the left chamber; Test2: a novel mouse introduced to the right chamber. Bottom: Representative movement traces from control and GluD1 KO mice across tests.

(B) Social index (SI) ratios were calculated by first normalizing each chamber to its respective prior test to control for chamber preference, followed by comparison of social vs. non-social targets within each test.

(C) Summary graph shows no change of SI ratios between control (n=12) and GluD1 KO (n=14) across Test1 and Test2.

(D-F) Similar results using the traditional method of measuring discrimination index for each test.

Unpaired Two-tailed t-test or Mann-Whitney U test revealed no significant differences. Data are mean ± s.e.m.

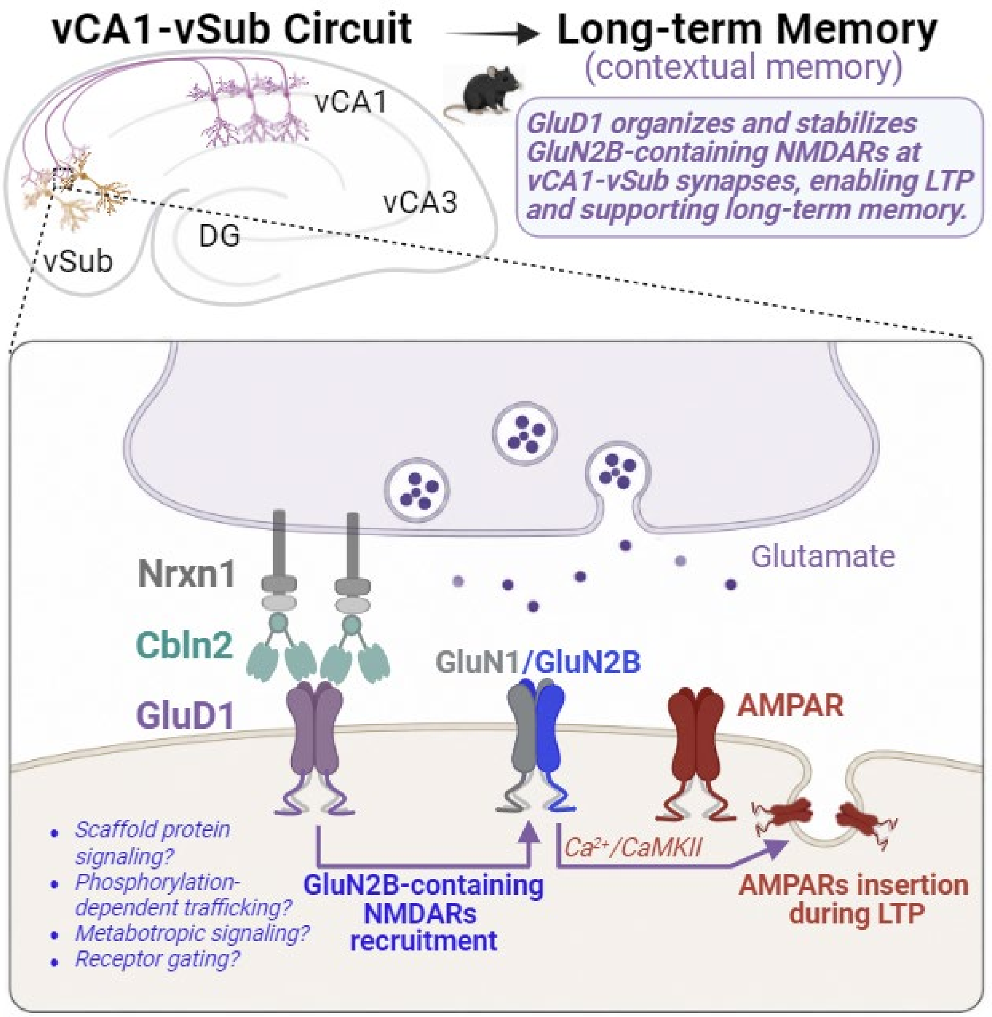


**Figure S5. Proposed model for GluD1-dependent regulation of GluN2B-containing NMDARs at vCA1→vSub synapses.**

The schematic illustrates the vCA1→vSub circuit and a working synaptic model in which the presynaptic Nrxn1-Cbln2 complex interacts with postsynaptic GluD1 to maintain GluN2B-containing NMDARs at vCA1→vSub synapses. GluD1-dependent stabilization or recruitment of GluN2B-containing NMDARs is proposed to support NMDAR-mediated $Ca^{2+}$ influx and downstream $Ca^{2+}$/CaMKII signaling, thereby promoting AMPAR insertion during long-term potentiation (LTP). Through this mechanism, GluD1-dependent GluN2B-NMDAR signaling supports synaptic plasticity and contextual long-term memory. The diagram also highlights several nonexclusive mechanisms that remain to be resolved, including effects on receptor trafficking and synaptic retention, postsynaptic scaffolding and signaling, phosphorylation-dependent regulation, metabotropic signaling, and potentially receptor gating.
